# The CTCF Anatomy of Topologically Associating Domains

**DOI:** 10.1101/746610

**Authors:** Luca Nanni, Cheng Wang, Freek Manders, Laszlo Groh, Paula Haro, Roel Oldenkamp, Stefano Ceri, Colin Logie

## Abstract

Topologically associated domains (TADs) are defined as regions of self-interaction. To date, it is unclear how to reconcile TAD structure with CTCF site orientation, which is known to coordinate chromatin loops anchored by Cohesin rings at convergent CTCF site pairs. We first approached this problem by 4C analysis of the *FKBP5* locus. This uncovered a CTCF loop encompassing *FKBP5* but not its entire TAD. However, adjacent CTCF sites were able to form ‘back-up’ loops and these were located at TAD boundaries. We then analysed the spatial distribution of CTCF patterns along the genome together with a boundary identity conservation ‘gradient’ obtained from primary blood cells. This revealed that divergent CTCF sites are enriched at boundaries and that convergent CTCF sites mark the interior of TADs. This conciliation of CTCF site orientation and TAD structure has deep implications for the further study and engineering of TADs and their boundaries.

## Introduction

It is generally accepted that transcription factors govern gene expression by recognizing specific DNA sequences, so-called transcription factor DNA motif instances, at gene promoters and enhancers. A major open question in this field is how enhancers, that are sometimes more than 100 kb from their target promoters, only stimulate one or a given set of gene promoters, rather than any other promoters they could come in contact with inside the nucleoplasm (Atlasi et al., 2019; Mifsud et al., 2015; Sanyal et al., 2012)?

Spatial segregation of chromosomal DNA segments would ‘shield’ genes from undesired *trans*- and *cis*-acting stimulatory interactions emanating from other transcription factor-bound chromosomal loci. Furthermore, spatial segregation of chromosomal loci may be coupled directly to the systems that compact chromatin domains into sub-micrometre volumes.

The biophysical phenomenology underlying storage of centimetre-sized DNA molecules into sub-micrometre volumes is a current knowledge frontier. The 23 human chromosomes harbour slightly less than 20,000 protein-coding genes, of which 3,541 exceed 100 kb, equivalent to a DNA length of 34 micron. Hence, two tasks nuclei must fulfil appear to oppose each other. Sub-micrometre chromatin compaction must be maintained while tens of micrometres of DNA must be reeled through the catalytic sites of RNA polymerase II in a regulated fashion so as to achieve proper gene expression levels.

Spatial DNA segregation implies the existence of boundaries along the length of chromosomes; namely DNA regions that insulate promoters from enhancers located beyond them. Moreover, isolation of a gene enhancer-promoter system within a chromosome requires two boundaries; one to the left and one to the right (Liu et al., 2015). In 1991 Kellum and Schedl (Kellum and Schedl, 1991) reported that flanking a transposon-borne transgene with the *Drosophila* heat shock 70 locus boundary elements can insulate it from chromosomal integration site position effects. Later, in 2001, an ill-explained phenomenon was reported; sometimes duplication of a boundary could nullify its insulating capacity (Cai and Shen, 2001; Muravyova et al., 2001). To this date, the molecular phenomenology that underlies chromatin domain boundary function is not fully understood.

As spatial segregation is a physical property, it can be investigated by slicing-up nuclei (Beagrie et al., 2017) or by using a molecular biology toolbox based on proximity-ligation of restriction enzyme-cleaved formaldehyde-fixed nuclear DNA (de Wit and de Laat, 2012; Dekker et al., 2013; Lieberman-Aiden et al., 2009).

Proximity ligation has revealed the existence of chromosomal topologically associated domains (TADs) of 10^5^ to 10^6^ bp in size (Nora et al., 2012). TADs are flanked by boundaries that are discovered as loci where there is a sharp break from preferential left-ward interactions to preferential right-ward interactions. In 2012, Dixon et al, thus modelled HiC data to detect 1,723 human TAD boundaries (Dixon et al., 2016; Dixon et al., 2012). Similarly, the Blueprint consortium found between 2,800 and 3,741 TADs in primary human blood cell types (Javierre et al., 2016). On the other hand it was shown that convergent CTCF sites are located at the synapses between long range interacting DNA regions, explaining multiple features of HiC data sets (Rao et al., 2014; Sanborn et al., 2015). Hence the current view is that large loops are constrained at their basis by Cohesin rings (Alipour and Marko, 2012; Dorsett, 2019; Fudenberg et al., 2016; Nasmyth, 2001; Parelho et al., 2008; Wendt et al., 2008) that accumulate at pairs of convergent CTCF sites to organize chromosome 3D architectures (de Wit et al., 2015; Dixon et al., 2016; Guo et al., 2015; Haarhuis et al., 2017; Nichols and Corces, 2015; Ong and Corces, 2014; Pugacheva et al., 2015; Rao et al., 2014; Remeseiro et al., 2016; Ruiz-Velasco et al., 2017; Sanborn et al., 2015; Vietri Rudan et al., 2015; Zuin et al., 2014).

There are reports of more than 200,000 accessible CTCF motif instances (CCGCGNGGNGGCAG) in the human genome (Liu et al., 2015) and Encode reported more than 60,000 ChIP seq peaks for CTCF in a large number of cell types (Encode Consortium, 2012). Not necessarily all these CTCF sites form loops, if one postulates that convergent CTCF sites are brought together by a (semi)processive extrusion-like process that will start by folding a chromatin strand on itself and then reel-in DNA to extend the nascent loop (Alipour and Marko, 2012; Nasmyth, 2001; Sanborn et al., 2015). Indeed, a single molecule DNA extrusion reaction has been reconstituted in vitro with Condensin, which is a distant structural relative of Cohesin (Hirano and Mitchison, 1994; Losada and Hirano, 2005), demonstrating ATP-dependent loop extrusion activity that can exert force on distal DNA tethers (Ganji et al., 2018).

To understand what makes up a functional boundary, we asked where the boundaries of the well-insulated glucocorticoid-inducible *FKBP5* gene are and what constitutes them. The results of this systematic exploration of the *FKBP5* locus were consistent with the hypothesis that TAD structure is hardwired by genomic CTCF site orientation. We then generalized our model by probing the topological relations between CTCF site orientations that could mechanistically explain TAD boundary function throughout the human genome. Altogether, our wet-lab experiments, bioinformatic findings and theoretical considerations provide a new formal CTCF site orientation classification with fundamental implications for the discovery and design of CTCF-dependent chromatin domain boundaries.

## Results

### 4C analysis of the chr6p21.31 locus

To explore topological organization and dynamics in and around the glucocorticoid-inducible *FKBP5* gene we deployed 15 ‘one-to-all’ 4C viewpoints (vp1-vp15, (van de Werken et al., 2012)). Altogether, the 4C viewpoints span 17 genes and 33 CTCF sites in 890 kb at chr6p21.31. The 4C results are plotted on a 1 Mb genome window, where viewpoints are arranged from left to right (Fig. 1). This window is equivalent to 0.34 millimetres of DNA that must be compacted length-wise at least a 1000-fold in order to fit inside a chromosome territory (Bolzer et al., 2005).

**Figure 1:**
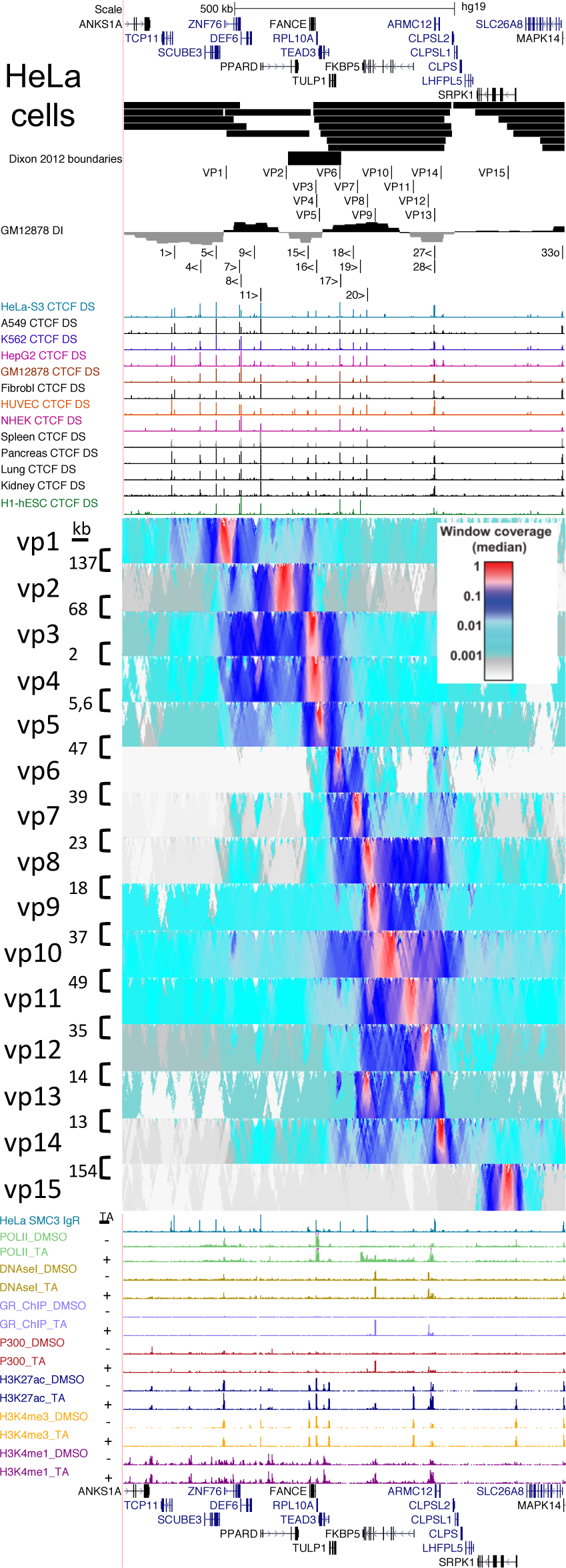
4C-seq walk across the *PPARD* and *FKPB5* genes and their TAD boundary at chr6p21.31. UCSC genome browser tracks display human genes, blood cell type TADs (Javierre et al., 2016), fifteen 4C viewpoints, the DI index of GM12878 lymphoblastoid cells and thirteen ENCODE CTCF ChIPseq tracks (Consortium, 2012), followed by the fifteen indicated 4C viewpoints from HeLa cells analysed using the 4C-seq pipeline (van de Werken et al., 2012). Below, the ENCODE HeLa Cohesin subunit SMC3 ChIPseq tracks is displayed and tracks for RNA polymerase II, DNaseI accessibility, glucocorticoid receptor, P300, H3K27ac, H3K4me3 and me1 from (Rao et al., 2011) are displayed, for samples treated or not with the synthetic glucocorticoid agonist triamcinolone acetonide (TA).

Two TADs are easily discerned on Figure 1, encompassing *PPARD* and *FKBP5*, which are long genes. The *PPARD* gene is located in a TAD detected as such by HiC by Javierre et al (Javierre et al., 2016) in CD4 and CD8 T-cells as well as being outlined by the TAD boundaries of neighbouring TADs detected in other blood cell types (Fig. 1). To the right of the *PPARD* TAD, the *FKBP5* gene is also located in a well-defined TAD that was detected in all analysed blood cell types. This indicates that 4C viewpoints are congruent with TADs defined by analysis of directionality index (DI) sign fluctuations (Dixon et al., 2012).

Between *PPARD* and *FKBP5* lies a 120 kb TAD boundary reported by Dixon et al in 2012 that separates the *FKBP5* TAD from the *PPARD* TAD. Five genes overlap this Dixon boundary (*PPARD*, *FANCE*, *RPL10A*, *TEAD3* and *TULP1*) as well as CTCF sites 13-17, although only CTCF sites 15, 16 and 17 show strong ChIPseq signal (Fig. 1). Viewpoints 1, 2 and 6 probe the edges of this boundary, and viewpoints 3, 4 and 5 explore the *RPL10A* promoter-associated CTCF16(<) site at the centre of this Dixon boundary.

### Insulation of PPARD by convergent CTCF sites

Close inspection of the 4C viewpoints on Figure 1 suggests that they may be classified in two types. Type I highlight one chromosomal region, examples are viewpoints 3, 4, 8-12 and 15. Type II highlight two or, more rarely, three regions. An extreme Type II interaction is displayed by viewpoint 13. This viewpoint shows more frequent interactions 153 kb further along the chromosome than locally at the viewpoint. Other Type II viewpoint examples are 1, 6, 7 and 14 (Fig. 1). The Type I/Type II viewpoint distinction indicates alternating interaction frequency regimes along the length of the chromosome. For instance, viewpoints 1 and 2 in the introns of *ZNF76* and *PPARD* display quasi anti-correlated interaction patterns, mutually ‘jumping’ across a stretch of chromosome they interact less with. In more detail; the first region highlighted by vp1 is at the left end of the *PPARD* TAD and spans 137 kb situated between the divergent CTCF4(<) and CTCF11(>) sites. This region encompasses *SCUBE3*, *ZNF76* and *DEF6* but not *PPARD*. The second region highlighted by vp1 is located beyond *PPARD* towards *FKBP5* and is rather small, spanning 19 kb located between CTCF15(<) and CTCF16(<). Hence vp1 highlights both edges of the *PPARD* TAD some 10-fold more than the inside of the *PPARD* TAD which is itself highlighted by vp2 as the 126 kb CTCF11(>) to CTCF(16<) interval.

These results support the notion that CTCF sites are organising principles for DNA-DNA interaction frequencies for this locus, as proposed at the global scale (de Wit et al., 2015; Haarhuis et al., 2017; Ong and Corces, 2014; Rao et al., 2014; Sanborn et al., 2015).

### 4C analysis of the RPL10A promoter CTCF site

*RPL10A* is the best expressed gene within 1 Mb and at the same time it is part of the left boundary of the *FKBP5* TAD. CTCF16(<) is located 0.3 kb from the TSS of *RPL10A*. We designed vp3, vp4 and vp5 around CTCF16(<). The viewpoints are located to the left of the nucleosome depleted, CTCF-bound *RPL10A* promoter 0.5 kb from CTCF16(<) (vp3), in the 4^th^ intron of *RPL10A* (vp4), and in the 12^th^ intron of *TEAD3* (vp5), being separated by respectively 2 and 5.6 kb.

Both vp3 and vp4 specifically highlight the convergent CTCF11(>) 126 kb to the left of CTCF16(<) with a spike of interactions in the same frequency regime as local proximity ligation around CTCF16(<) (red signal, Fig. 1). However, while to the left vp3 homogenously interacts up to and beyond CTCF(11>), the vp4 viewpoint shows a slight dip in its interaction profile in the middle of the *PPARD* TAD, where vp2 is located. Following this trend, vp5 which is only 5.6 kb to the right of vp4, shows more of a Type II interaction pattern that is rather complementary to the *ZNF76* vp1 type II pattern. Viewpoints 3, 4 ad 5 highlight the same intervals flanking the primary *PPARD* loop. To the left, their interactome extends beyond CTCF11(>), highlighting a region of 100 kb, almost up to CTCF5(<), beyond CTCF 7(<) and CTCF8(>) which are only 3 kb apart. To the right, the interactome of vp3, 4 and 5 appears to be delimited by CTCF17(>).

It could therefore be argued that the three viewpoints at the RPL10A promoter CTCF16(<) site highlight left and right *PPARD* ‘sub-TADs’ that consist of DNA up to secondary ‘back-up’ CTCF sites flanking the primary CTCF11(>) - CTCF16(<) *PPARD* 126 kb loop, in a <><><<> pattern of CTCFs 5, 7, 8, 11, 15, 16 and 17 that spans 281 kb and appears to constitute the entire *PPARD* TAD.

### The PPARD-FKBP5 boundary

Viewpoint 6 is located 0.5 kb to the left of CTCF17(>), which in turn is 29 kb to the left of CTCF18(<). Viewpoint 7 is 39 kb to the right of vp6, beyond CTCF18(<) and 7.5 kb left of CTCF19(>). Both vp6 and vp7 highlight the same local region of about 110 kb that flank the Dixon boundary between the *FKBP5* and *PPARD* TADs. The left limit of this region appears not to be a CTCF site, it is the bivalent H3K27me3 promoter of *TEAD3* gene located between CTCF16(<) and CTCF 17(>) inside the Dixon boundary. This limit is shared by vp6, 7, 8, 10 and 14 (Fig. 1), indicating its recurrent character as a limit for viewpoint interactions. The right limit of the local interactome of vp6 and vp7 is the CTCF20(>) site. The region between viewpoints 5, 6 and 7 is a *bona fide* boundary between the *PPARD* and *FKBP5* TADs, in that vp5, located 6 kb to the right of CTCF16(<) shows Type II far-left interactions but 47 kb further to its right, vp6 at CTCF17(>) displays Type II far-right interaction. Following the ‘mirroring trend’ set by vp1 and vp5, vp7 is mirrored by Type II viewpoints 13 and 14. Viewpoints 7 and 14 thus appear to be excluded from the *FKBP5* gene loop much like vp1 and vp5 are excluded from the *PPARD* gene loop. The CTCF architecture of the sub-TAD highlighted by vp 6 and 7 ends at the bivalent *TEAD3* promoter and consists of CTCF17(>), CTCF18(<), CTCF19(>) and CTCF20(>), which is followed by the primary loop of the *FKBP5* TAD whose right-side anchor is the closely spaced CTCF27/28(<<) pair. This yields an *FKBP5* TAD CTCF17-28 pattern that can be written as ><>> <<, if one only considers CTCF sites that yield high occupancy levels by ChIPseq (Fig. 1, (Encode Consortium, 2012)).

In summary, the 4C results suggest that the boundary between *PPARD* and *FKBP5* harbours two sub-TADs that are revealed by Type II viewpoints. One sub-TAD belongs to the *PPARD* TAD and the other to the *FKBP5* TAD. Between these sub-TADS, the H3K27me3/H3K4me3-marked bivalent *TEAD3* promoter (Bernstein et al., 2006) located between CTCF16(<) and CTCF 17(>) appears to be the physical interaction border of 4C viewpoints.

### FKBP5 TAD structure is resilient to increased transcription rates

We took advantage of the fact that glucocorticoids stimulate *FKBP5* transcription 15 to 50-fold in almost all human cell types to investigate the relation between transcription rates and locus architecture (Fig. 1, bottom legend, Fig. S1A).

Viewpoint 8 is located 0.8 kb to the right of CTCF20(>) in intron 8 of *FKBP5* (Fig. 1). It behaves much like vp3 and vp4 for the *PPARD* TAD, as it contacts the entire *FKBP5* TAD and shows a spike of interaction frequency in the same regime as local proximity ligation at CTCF27(<), located 153 kb away, beyond the steroid-inducible *FKBP5* super-enhancer at −29 to −45 kb relative to the *FKBP5* promoter. The limits of the CTCF20(>)-proximal vp8 interactome appears to be the bivalent H3K27me3 *TEAD3-TULP1* promoter in the 55 kb divergent DNA interval, bounded by CTCF16(<) and 17(>) at the heart of the Dixon boundary to the left, and the *CLPS1* heterochromatin region beyond *ARMC12* to the right. Intriguingly, but consistent with previous reports (D’Ippolito et al., 2018; Stavreva et al., 2015; Wang et al., 2019), glucocorticoid stimulation does not visibly affect vp8 *cis*-associations measured by 4C (Fig. S1A).

Viewpoints 9, 10, 11 and 12 respectively probe the glucocorticoid-inducible enhancer in the 5^th^ intron of *FKBP5*, a H3K4me1 region located in *FKBP5*’s 1^st^ intron, the promoter, and the −29 to −45 super-enhancer of *FKBP5*. The four viewpoints highlight the 153 kb loop that is anchored at CTCF20> and CTCF27(<), sometimes also highlighting the CTCF17(>) – CTCF20(>) interval (Fig. 1). None of these viewpoints are greatly affected by glucocorticoid stimulation of *FKBP5* transcription rates (Fig. S1A).

We employed viewpoints 8, 9, 10, 11 and 13 in purified human monocytes and found the same TAD architecture at *FKBP5* as in HeLa cells (Fig. S1B), indicating the existence of very similar chromatin interaction frequency domains in this non-proliferating primary human blood cell type.

### The CTCF27/28 right-hand boundary of the *FKBP5* TAD at the *ARMC12* promoter responds to increased transcription rates

To the right of *FKBP5* lies *ARMC12*, which is expressed only in testis. The *ARMC12* promoter lies between two CTCF sites, 27(<) and 28(<), which are separated by only 1.2 kb. Viewpoint 13 is located between CTCF27 and 28. Viewpoint 13 is the most extreme Type II viewpoint we encountered in that it harvests more reads 153 kb away around CTCF20(>) than locally at CTCF27/28(<<) (Fig. 1, S1A-C). Surprisingly, when transcription activity is low, the quasi-exclusive interaction of CTCF27/28(<) with CTCF20(>) is not reciprocated by CTCF20(>) (vp8, see above). This indicates that the 1.2 kb interval between CTCF27(<) and CTCF28(<) has particular topological associations. Interestingly, when transcription rates increase, CTCF27/28(<<) appears to sometimes detach from CTCF20 and attach to CTCF17(>) more often (Fig. S1A). We could replicate *FKBP5* transcription-induced increased pairing of CTCF17(>) with CTCF27/28(<<) in two other cell types; the established THP-1 monocytic leukaemia cell line and primary human blood monocytes of healthy blood donors (Fig. S1C). However, these transcription rate effects are not crisp as CTCF17(>) – CTCF27-28(<<) interactions were variably highlighted also under non-induced conditions, perhaps because *FKBP5* is always transcribed at moderate levels (range: 2 - 10 RPKM), with surges of transcriptional activity induced by steroids (range 30 - 150 RPKM).

Why do vp6 and vp13 report transcription rate changes? We speculate that this is because CTCF20(>) is located in intron 8 of *FKBP5*, 26 kb 5’ of the major polyadenylation site of the *FKBP5*(-) gene. The increased passage of the RNA polymerase II transcription bubble necessarily disrupts DNA double strand topology and thus, most likely, also paired CTCF site chromatin structures. RNA polymerase thus likely causes detachment of CTCF20(>, vp8) from CTCF27/28(<<, vp13), which now joins CTCF17(>, vp6) more often (Fig.S1A), presumably through chromatin extrusion processes. Furthermore, in the Hela and THP-1 cell lines as well as in primary monocytes, at high *FKBP5* transcription rates, CTCF27/28(<<) shows somewhat increased interactions with the *FKBP5* promoter and enhancers (Fig. S1C, vp13).

We conclude that although transcription rates can subtly affect *FKBP5* TAD architecture when RNA polymerase II passage disrupts CTCF-DNA interactions, the general architecture and the specific limits of spatial interaction frequency regime domains is maintained also in the face of increased transcription (Fig. S1D). This suggests that the hypothetical CTCF-bounded chromatin extrusion system can operate on time scales that are similar or faster than the time scales of chromatin loop disruption imposed by gene transcription.

### CTCF site spatial distribution analysis reveals orientation biases

In order to dissect the mechanistic roles of CTCF sites and their orientation in a genome-wide fashion, we started with the collection of 61,079 human CTCF sites reported by Rao et al (2014) and 33 ENCODE CTCF ChIPseq experiments (Table S1, (Encode Consortium, 2012)). More than 16,000 CTCF sites are occupied in the 33 samples and more than 40,000 are CTCF-bound in more than 20 samples.

The genome-wide median distance between CTCF sites is 21.6 kb (Fig. 2A), but the distribution of distances between CTCF sites is not uniform. CTCF-CTCF distances are smaller than expected in the 10^3^-10^5^ bp range (Fig. 2B).

**Figure 2:**
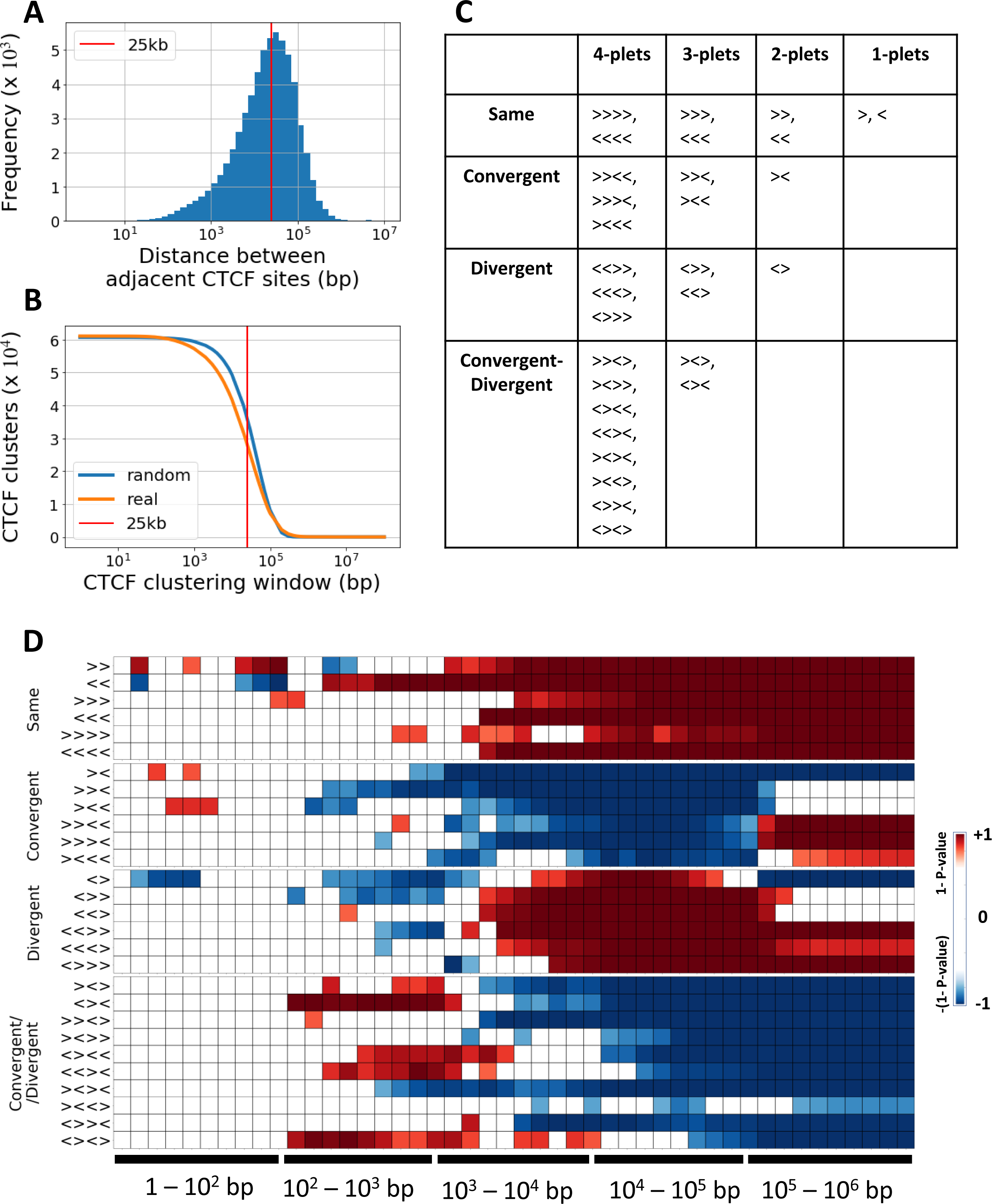
CTCF sites spatial distribution analysis reveals orientation biases. A. Distribution of distances between CTCF binding sites along the human genome. The 25kb distance is indicated as a red vertical bar. B. Number of CTCF clusters at varying clustering window. Starting with a window of 1 bp yields 61,079 mono-plets and using a 10^8^ window yields 46 clusters (one per chromosome arm). Shuffled CTCF sites along the genome (blue) are compared to the real spatial distribution of sites (orange). The red line shows the 25 kb clustering window. C. Theoretically possible relative orientation combinations for one to four CTCF sites. They belong to classes *Same* (all sites oriented in the same direction), *Convergent* (sites pointing towards each other), *Divergent* (sites pointing away from each other) and, for three and four CTCF sites, the class *Convergent + Divergent*. D. Statistical assessment of over-(red) and under-represented (blue) occurrences of CTCF patterns along the indicated window ranges. The five indicated window ranges are divided in ten equal bins. P-values are from binomial tests for each pattern and bin separately for diplets (1/4), tri-plets (1/8) and tetra-plets (1/16). Patterns are arranged according to panel (C). The colour scale reflects the p-values resulting from the two one-sided tests.

We hypothesized that same, convergent and divergent CTCF pairs would not show the same 1D spatial distribution along the length of chromosomes. Therefore, we investigated CTCF site clusters as a function of their relative orientations (Fig. 2C). For mono-plets there are left (>) and right (<) orientations. For di-plets there are two patterns oriented in the same direction, one convergent and one divergent. For tri-plets there are two same-oriented, two convergent + same oriented, two divergent + same and two convergent + divergent patterns. For tetra-plets there are 16 patterns; two same oriented, three convergent, three divergent and eight convergent + divergent. Altogether, CTCF site permutations of 3 or more sites can therefore by divided into 4 distinct classes, depending on the occurrence or not of convergent and divergent CTCF pairs (Fig. 2C, see Methods).

Next, we plotted the spatial distributions of each pattern as a function of increasing genomic window sizes and determined the deviations from the expected binomial distributions (Fig. 2D). Given a window *w*, we counted *n*-plets (with *n* from 2 to 4) whose maximal CTCF-CTCF site distance is *w* or less. We then compared this number to the expected value (1/2^n^) for each pattern using the binomial test (See Methods).

This revealed that for any *n*-plet class, same, convergent, divergent and convergent + divergent CTCF blocks yield very similar spatial distribution patterns, which are different for each class (Fig. 2D).

Same patterns tend to be more prevalent in 1 kb windows onwards. Convergent patterns are more prevalent in windows exceeding 200 kb and depleted from 3 to 100 kb windows. By contrast, divergent patterns are more prevalent from 5 to 100 kb and depleted from 0.3 to 2 kb. Finally, convergent + divergent CTCF patterns show a mixed picture but are still more prevalent from up to 0.1 kb to 3 kb while they tend to be depleted from 100 kb and larger windows (Fig. 2D).

To test the robustness of this result we associated an aggregate rank score to each CTCF site by multiplying their ChIPseq signal and CTCF motif score ranks (See Methods). The aggregated rank score was used to partition CTCF sites into 4 quartiles (Fig. S2G) which were successively removed prior to re-computation of spatial CTCF distribution density maps. This revealed that the enrichment of CTCF patterns in different window sizes was robust to removal of the lowest 25% of sites (Fig. S2B). Retaining only the top 50% of sites largely preserved the spatial density imbalances except for long range same CTCF patterns which became depleted (Fig. S2D). Strikingly, convergent di-plets (><) which are depleted when using the entire CTCF set are enriched when only the top quartile is retained (Fig. S2F), suggesting that CTCF sites constituting long range (10^5^ to 10^6^ kb) convergent pairs have high rank. Similarly, removal of the top 25% of CTCF sites also preserves the prevalence of divergent CTCF patterns in the 20 to 300 kb range (Fig. S2E).

We conclude that CTCF site arrangements are not random. Convergent and divergent CTCF sites tend to show complementary spatial densities as a function of genomic distance. Convergent CTCF sites being more prevalent at long distances (> 100 kb) while divergent CTCF sites are more prevalent at intermediate distances (<100 kb). Moreover, these properties are preserved when low ranking CTCF binding sites are removed from the analysis.

### Generating a gradient of boundary identity

In order to dissect the mechanistic roles of CTCF sites at boundaries we needed to identify *bona fide* boundaries. Hereto we searched for topological boundaries that are preserved in multiple human blood cell types, namely the HiC TAD boundaries reported by the Blueprint consortium (Javierre et al., 2016) for seven adult human blood cell types. The TADs ultimately called by Javierre et al were coherent in two biological replicates. To find consensus boundaries we developed a ‘boundary consensus algorithm’. The algorithm takes as input a set of boundary sets and outputs a single set of boundaries together with a conservation score *s* that indicates in how many cell types the boundary was called. Fig. 3A visually details this procedure (See Methods).

**Figure 3:**
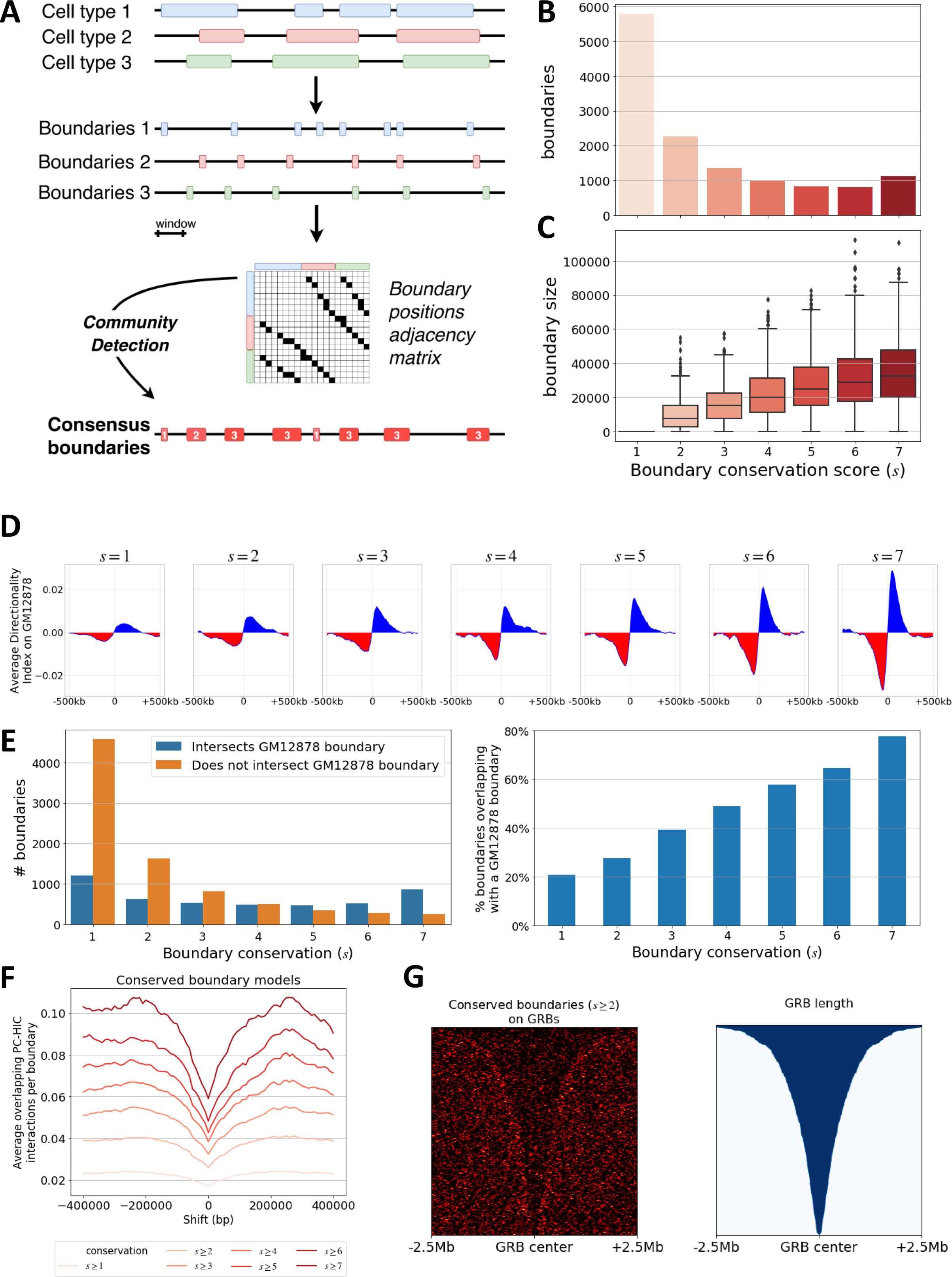
Conserved boundaries show a gradient of enrichment of HiC features. A. Schematic representation of the boundary consensus algorithm using an example of three cell types and ten TADs to yield eight boundaries with conservation scores 1 to 3. B. Number of detected boundaries as a function of conservation score. C. Boundaries size distributions as a function of conservation score. D. Average Directionality index in 5 kb computed on GM12878 and projected onto s1 to s7 boundaries in 1 Mb windows. E. Number (left panel) and percentage (right panel) of s1-s7 boundaries that intersect a GM12878 boundary. F. Percentage of PC-HIC interactions with a CHICAGO score greater than 5 that span a consensus boundary. At the centre the real boundary set is shown and then the coordinates are shifted left and right in windows of 10 kb. This metric is plotted for every consensus boundary set from *s* ≥ 1 to *s* ≥ 7. G. Density of consensus boundaries on GRBs aligned at the centre of 5 Mb regions and ordered from larg to small.

The algorithm detects 13,131 boundaries of which 5,786 are ‘cell-type-specific boundaries’ in that they have a conservation score *s* of 1. The results we obtained suggest that the boundary consensus algorithm is starting to saturate, since the number of boundaries with a conservation score of 4, 5 or 6 are individually less numerous than the boundaries detected in 7 cell types (s7, Fig 3B). The s2–s7 consensus boundaries occupy 152 Mb of the genome. The most conserved are larger as they are derived from the largest number of cell types (Fig. 3C). They have a median size of 32.5 kb.

To independently validate the boundaries, we included GM12878 cell line HiC data (Rao et al., 2014) in our analysis. The site of sign inversion from a negative to a positive DI is the landmark to call HiC boundaries (Dixon et al., 2012). Figure 3D shows the DI of GM12878 cells centred on the boundaries of each conservation level (s1-s7). This reveals a quasi-linear increase in the sharpness of the boundaries as a function of their conservation (Fig. 3D), indicating that as conservation increases, the DI increases, too. It is therefore likely that many of the consensus boundaries are also boundaries in GM12878. We verified this by intersecting the 6,073 boundaries of GM12878 cells with our boundaries (s1-s7). Overall 20% of the s1 boundaries intersect a GM12878 boundary and this ratio increases linearly up to 80% of the s7 boundaries (Fig. 3E). This indicates that highly conserved boundaries have a high chance of existing in other cell types.

To assess boundary function in gene expression, we took the approach of Schoenfelder et al (Schoenfelder et al., 2015), and generated ‘insulation plots’ whereby promoter capture HiC (PC-HiC) revealed 723,600 gene-promoter interactions in the seven blood cell type data of Javierre et al (Javierre et al., 2016) with a CHICAGO score greater than 5. The result (Fig. 3F) shows that the boundaries’ insulation capacity is positively correlated with their consensus score (See Methods).

As a final independent confirmation of boundary function, we projected the s2-s7 boundaries on a set of 815 Genomic Regulatory Blocks (GRB, (Harmston et al., 2017)) conserved throughout vertebrate evolution. The consensus boundaries clearly delineate many of these regions, although some are interrupted by boundaries, possibly indicating evolutionarily conserved gene regulatory architectures that involve two or more adjacent TADs (Fig. 3G).

The above results demonstrate that it is possible to stratify boundaries based on their conservation across blood cell types and that features derived from external HiC datasets and regulatory interactions can be quantitatively assessed as a function of this stratification gradient.

### Conciliation of consensus boundaries and CTCF sites reveals divergent CTCF sites at boundaries

With the spatial distribution of CTCF orientation pattern classes along the genome and a set of boundaries with increasing boundary identity in hand, we wished to ask which CTCF patterns code for TAD boundary function.

Intersecting the 7,345 s2-s7 consensus boundaries with CTCF site locations revealed that 16% lack a CTCF site even when we scanned an additional 25 kb to the left and to the right of the boundary’s borders. To gain more insight, we projected the 3,388 CTCF-less boundaries on our gradient of boundary identity. The proportion of boundaries without CTCF sites decreases as boundary conservation increases (Fig. 4A). About half of the CTCF-lacking boundaries are conserved one or two times. This suggests that a small proportion of boundaries might perhaps exert their function without CTCF, but this would concern a minority of the highly conserved boundaries.

**Figure 4:**
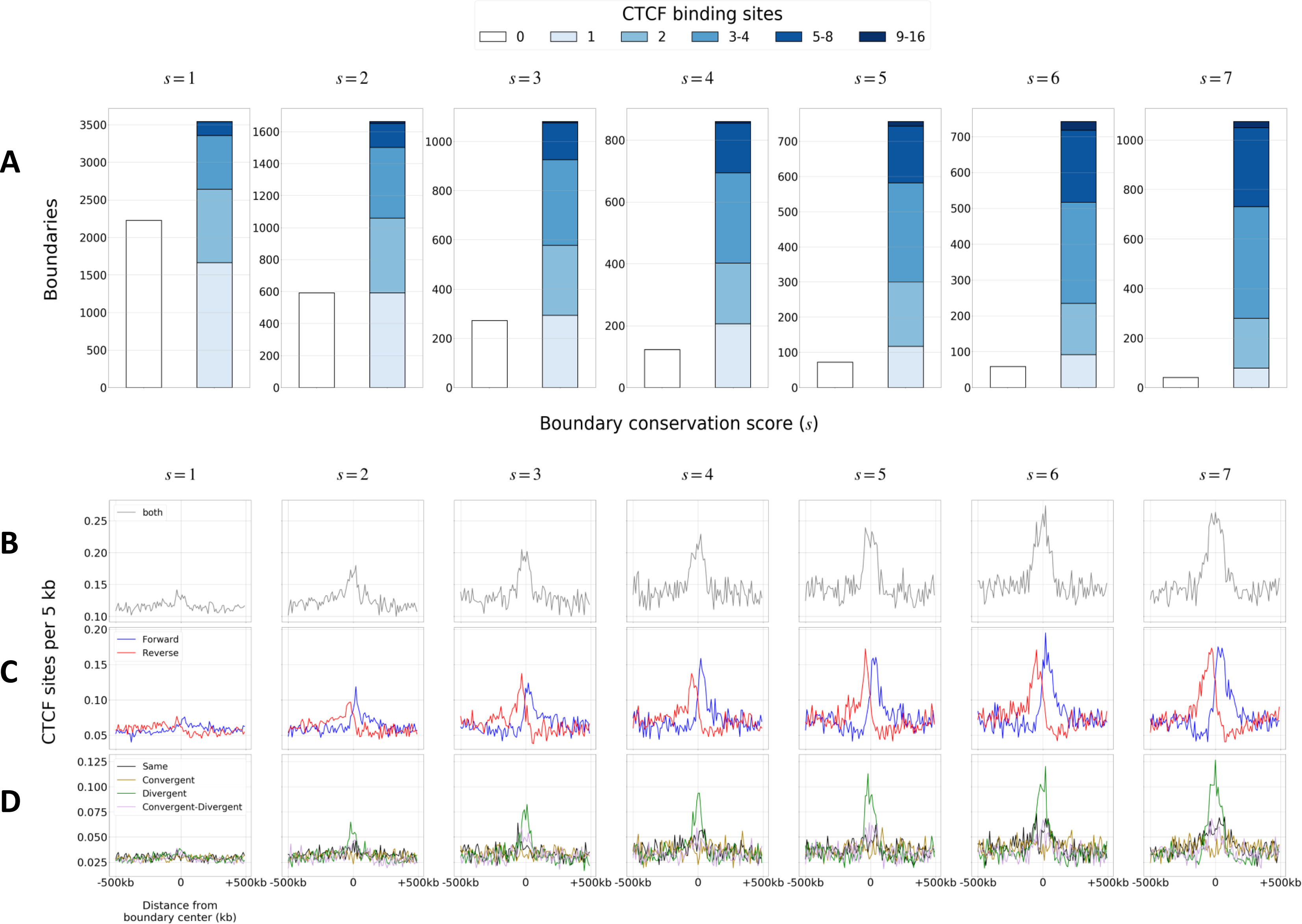
Consensus boundaries are enriched of divergent CTCF sites. A. Number of boundaries with a given conservation score that harbour 0 to 16 CTCF sites. B. Average number of CTCF sites per 5 kb in a 500 kb window around the boundary centres. C. Average number of CTCF sites per 5 kb in a 500 kb window stratified on their > and < orientation. D. The CTCF triplet pattern naming convention shown on Fig. 2C was used to classify CTCF binding sites according to their two adjacent CTCF sites’ orientations.

To obtain insight into mechanistic roles of CTCF site orientations at boundaries we rendered CTCF site density on 500 kb regions centred on the middle of the boundaries (See Methods). As expected, this revealed enrichment of CTCF sites at boundaries (Fig. 4B). Next, we partitioned CTCF sites into ‘Reverse’ (<) and ‘Forward’ (>) CTCF sites (Fig. 4C). This split the signal observed on panel 4B into two sets, peaking to the left and to the right of the boundaries, demonstrating that reverse CTCF sites (<) tend to locate to the left side of boundaries while forward CTCF sites (>) tend to locate to the right side. Next, we assigned a class to each of the 61,079 CTCF sites following the triplet nomenclature (same, convergent, divergent or convergent + divergent, see Fig. 2C) that uniquely defines each CTCF site as a function of its left and right neighbours’ orientations (See Methods).

Remarkably, only one of the CTCF orientation classes shown on Fig. 2C is enriched at boundaries, namely the divergent class. This observation is buttressed by the fact that enrichment of divergent CTCF sites is proportional to the boundary consolidation score, closely following the DI index increase plotted on Fig. 3C. Hence, the more a boundary is easy to detect, the more likely it is to harbour at least one divergent CTCF site.

These results suggest that ‘boundary function’ is mechanistically conferred by divergent CTCF sites, namely <<> and <>>.

### The CTCF anatomy of TADs

The definition of a boundary is a DI sign change from negative to positive, indicating a point where DNA interaction frequency biases sharply shift from a bias to the left to a bias to the right (Fig. 1A, (Dixon et al., 2015)). A logical corollary of this is that there will be the opposite sign change inside TADs so as to switch from left-biased interactions to right-biased interactions abutting the next boundary.

Where precisely in a TAD this (+) to (-) DI sign inversion will take place may be determined biophysically through chromatin fibre dynamics and genetically though CTCF site orientation positions (Racko et al., 2018).

We first investigated the possibility that CTCF sites determine TAD’s (+) to (-) sign inversion regions by building a ‘meta-TAD’ that represents the entire collection of 7,435 TADs as an equal number of bins whose real size depends on TAD length. This reveals a cline of Forward (>) CTCF sites on the left half of the meta-TAD. Conversely, the right half shows a cline of Reverse (<) CTCF sites (Fig. 5A-C).

**Figure 5:**
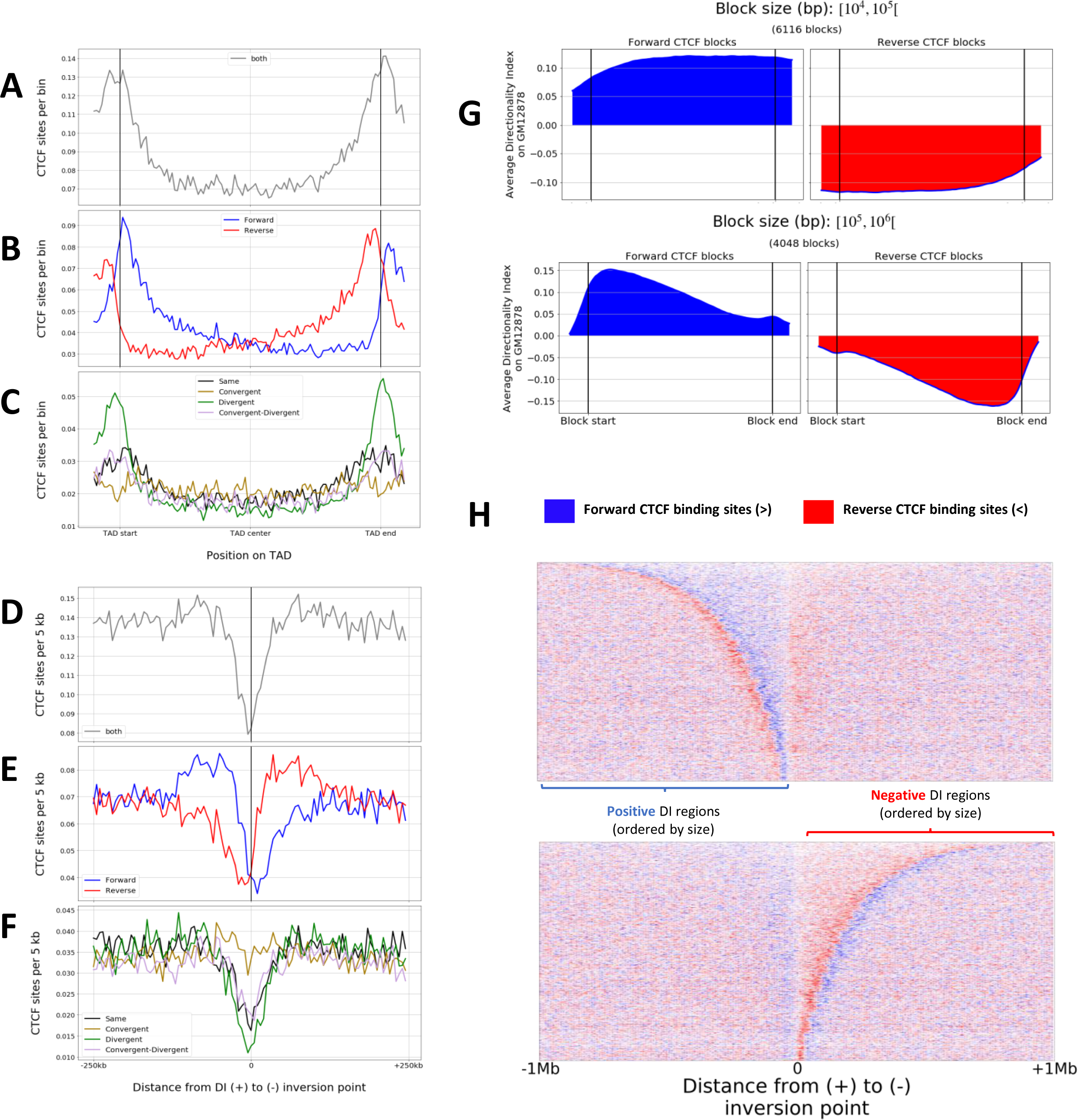
TAD centres are depleted of all CTCF patterns except convergent pairs. A-C Meta-TAD representation to show the enrichment of CTCF binding sites (A), CTCF site orientation (B) and CTCF orientation classes (C). Each TAD was divided in 100 bins and the average number of CTCF sites was computed for each bin. D-F 500 kb windows around the (+) to (-) points of inversion of DI index showing per 5 kb the average number of CTCF sites (D), CTCF site orientation (E) and CTCF orientation classes (F). G Average DI index on arrays of same-oriented CTCF sites in Forward (left panel) and Reverse (right panel) orientation ranging in size from 10^4^-10^5^ (top) or 10^5^-10^6^ (bottom) bp. H Distribution of Forward (>, blue) and Reverse (<, red) CTCF sites in a 1 Mb window around the (+) to (-) points of inversion of DI index ordered by the length of the DI-positive (top) and DI-negative (bottom) section of the TAD.

This suggests that the ‘middle of the TAD’ location defined as a DI sign inversion from (+) to (-) coincides with a reversal in bias of CTCF site orientations.

As a first test, we calculated the average DI index on arrays of same-oriented Forward or Reverse CTCF sites of various length. In keeping with the hypothesis, we found that Forward and Reverse CTCF site regions are enriched in regions of positive and negative DI index, respectively (Fig. 5G).

To further investigate this, we inspected the regions around sites of (+) to (-) DI sign inversion and plotted CTCF site density. This revealed that sites of (+) to (-) DI sign inversion are depleted of same, divergent, and convergent + divergent CTCF sites. However, the density of convergent CTCF sites is preserved at 0.035 per 5 kb. Operationally, this results in a local enrichment of convergent CTCF (><<, >><) at the ‘middle’ of TADs whilst maintaining a ‘CTCF-depleted’ local environment (Fig. 5D-F).

Next, we wished to assess CTCF orientation bias inside the TAD halves with a positive versus those with a negative DI index. To this end, we isolated positive DI(+) regions, ranked them from longest to shortest and coloured 10 kb bins as blue (Forward CTCF >), red (Reverse CTCF <) or white when no CTCF site occurs in that bin. This revealed that the CTCF sites on either side of the (+) to (-) DI sign inversion are indeed more often of the same orientation as the divergent CTCF sites that mark the closest boundary, and that this obeys the DI sign (Fig. 5H). Boundaries thus appear to be safeguarded from ‘invasion’ by extrusion complexes originating inside TADs by sets of convergent CTCF sites that make up primary TAD loops that can be extended with sub-TADs.

We therefore speculate that the (+) to (-) DI sign inversion site between boundaries is partly hardwired through evolutionary preservation of convergent CTCF sites and depletion of other CTCF orientation classes. Hence, TAD CTCF anatomy seamlessly joins boundary CTCF anatomy as boundaries are enriched in divergent CTCF sites that point towards the bordering TADs interiors.

### Modulation of CTCF loop network geometrical constraints

An intriguing feature that Rao et al extracted from HiC data are the ‘Hi-C Computational Unbiased Peak Search’ (HiCCUPS) and their anchors (Rao et al., 2014). These anchors represent regions of distal interaction between regions whose flanking regions do not interact as frequently, yielding bright pixels in HiC contact maps. Operationally they are considered to be long range chromatin loops. Globally, 9,448 HiCCUPS loops were detected in GM12878 cells, of which 4,654 harbour a convergent pair of CTCF binding sites and Cohesin at their anchors (Rao et al., 2014).

We simulated the HiCCUP topological structure of the genome using the same logic as Sanborn et al (Sanborn et al., 2015). The simulations involved 100 epochs in which one extrusion event happened independently between each adjacent pair of CTCF sites.

We first evaluated the ‘full stop’ CTCF model, in which each site has minimum permeability (stopping probability equal to 1). The full stop model identifies 20% of the GM12878 HiCCUPS (Fig. 6A). This is due to the impossibility to create loops that span a smaller convergent loop. We therefore attempted to recall long HiCCUPS by parametrization of CTCF binding sites using motif and ChIPseq scores. To this end we assigned a *permeability* score to each CTCF site that is inversely proportional to its rank score (Fig. 2, see Methods). We then performed three more looping simulations: one using only motif score, one using only ChIPseq score and one with the rank aggregated score we had used to analyse the resilience of CTCF site class spatial distributions to CTCF site removal (Fig. 2D). At the genome-wide level, the rank aggregated parametrization greatly outperformed the other simulations in the recovery of HiCCUPS loops, both in terms of recall (Fig. 6A) and of precision (Fig. 6B). Our results therefore suggest that both motif strength and ChIPseq signal should be considered when simulating chromatin extrusion outcomes in terms of loops because the relation between these two parameters is variable.

**Figure 6:**
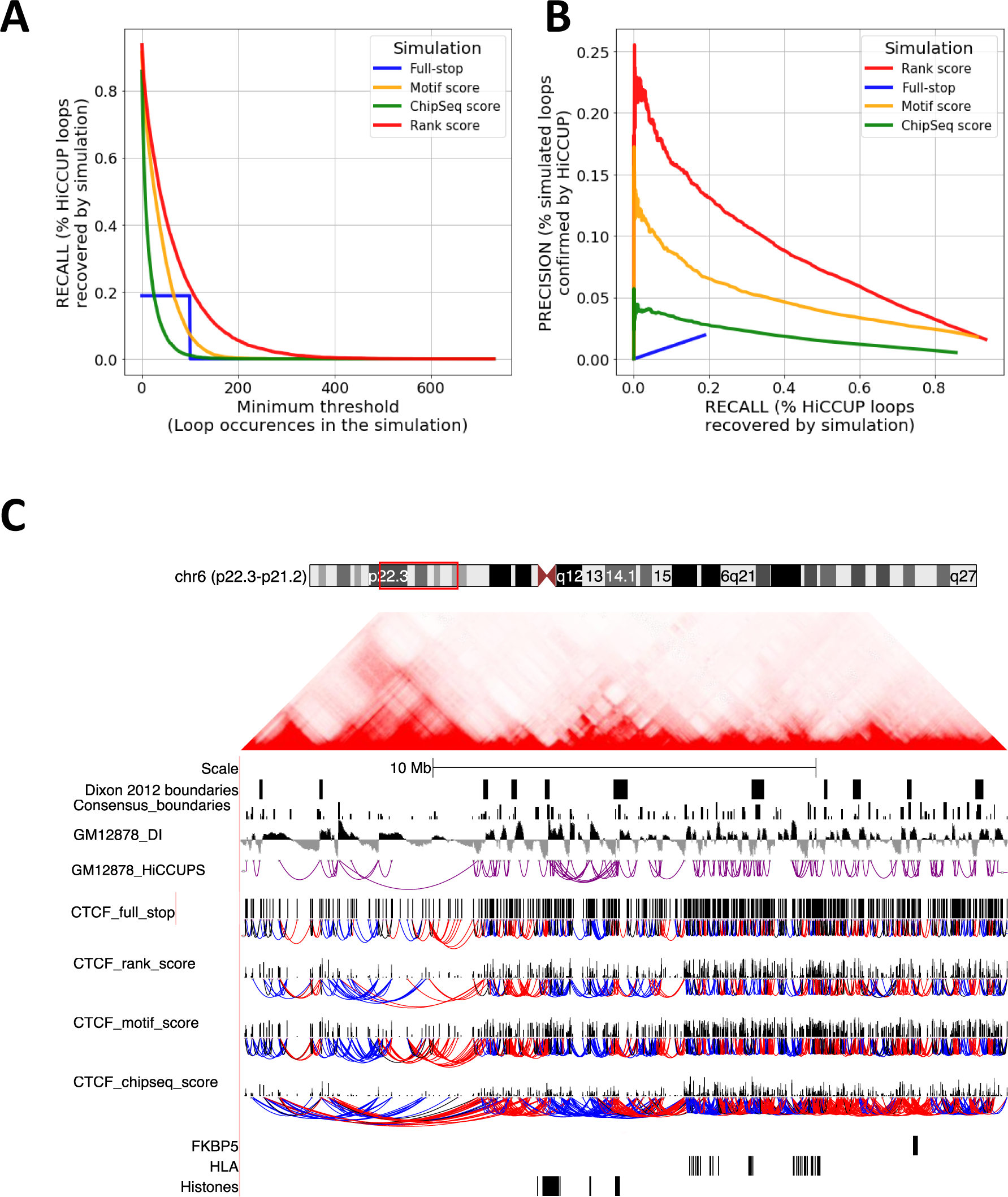
CTCF looping simulations work best when both ChIPseq and Motif scores are integrated. A Percentage of HiCCUPS loop (recall) that are recovered by the various models as a function of their occurrence in the simulations. B Precision-recall curves of the indicated models, indicating the percentage of simulated loops which are present in the HiCCUPS collection as a function of recall. C Loop simulation results in a 20 Mb window (chr6:18,000,000-,380,000,000) rendered on the UCSC browser (Kent et al., 2002) together with HiC data from GM12878 (50 kb resolution, Genome-wide balanced) rendered in juicebox (Durand et al., 2016a). Boundaries, DI index and HiCCUPS loops are indicated above the CTCF sites (Rao et al., 2014). Four indicated simulations and the associated CTCF permeability scores are shown. Loops are coloured according to the bias in orientation of the intervening CTCF sites between loop anchors. Blue is Forward (>) and red is Reverse (<) orientation-biased. The locations of the *FKBP5* gene, the MHCI and II (*HLA*) and the *HIST1* histone clusters are indicated below.

We investigated the potential for altered relations between CTCF motif scores and CTCF ChIPseq signal as a function of distance from the 5,886 gene transcription start sites (TSS) equipped with a potentially functional CTCF site such as CTCF16(<) at *RPL10A* (vp3-5, Figure 1). The lowest motif/ChIPseq score ratio is achieved within a 300 bp window 5’ of TSSs, indicating efficient CTCF binding at these sites, presumably because promoter-associated ATP-dependent chromatin remodelling machines facilitates access of CTCF to promoter-DNA binding sites (de Dieuleveult et al., 2016). The smoothened average ratio then increases to double its value between the TSS and 150 bp downstream and is maintained downstream of the TSS when highly expressed genes are considered (Fig. S3). We speculate that RNA polymerase II complexes compete there with CTCF for DNA binding. These results indicate that CTCF binding can indeed be modulated.

Interestingly, when we coloured simulated loops as a function of the underlying number of Forward and Reverse CTCF sites, roughly alternating patterns emerged, that can be schematised as <>…>≤ and ≥<…<>, whereby the ultimate convergent CTCF site (underlined) is a ‘recipient’ of multiple potential right or left CTCF interactions (Fig. 6C). Although the alternation of left and right array is inherent to any two-symbol grammar, for DNA, the permeability property simultaneously modulates the length (bp) of loops as well as the frequency of loop occurrence (time). This suggests that TADs should not be considered as static objects, but rather as dynamic topological entities bounded by active divergent CTCF sites that can form families of loops between convergent CTCF site pairs.

Finally, we note that in apparent contradiction to the general <>(>…>)(<…<)<> CTCF TAD model (Fig. 5H), both DI index-based and loop simulation results revealed that many individual left halves of TADs do harbour left pointing same CTCF arrays and vice versa. We speculate that this reflects evolutionary pressure to preserve locus-specific topologies that assist long-distance gene expression control via distal enhancers in the face of the evolutionary history of chromosomes during which inversions, deletions, duplications and translocations invariably took place.

### In Epigenomic signalling by the glucocorticoid receptor conforms to half TAD dimensions

In light of our new insights we hypothesized that long range enhancer-signalling might be confined to TAD halves. To this end we choose 401 robust glucocorticoid receptor (GR) binding sites detected by ChIPseq in purified primary human blood monocytes that induce 1,328 H3K27ac peaks in the TADs they reside (Wang et al., 2019). These reside in 374 half-TADs.

Of 13,161 TAD halves determined by DI index inversions, 235 harbour at least one GR peak and have four adjacent TAD halves without a GR peak. Up-regulated H3K27ac signal is strongly enriched in the GR-bearing TAD halves and there is also a small enrichment of up-regulated features, in the *n + 1* half-TADs with a (+) DI and in *n – 1* half-TADs for (-) DI regions with GR peaks (Fig. S4).

The above results suggest a model where CTCF sites and the TAD sub-structures they constrain are indeed the structural backdrop within which long-range transcriptional *cis*-acting regulatory interactions instigated by DNA-bound glucocorticoid receptors take place.

### Intragenic TAD boundaries and CTCF site orientations

Healthy blood donor monocyte-derived macrophages express 7,474 genes at 1-10 RPKM, 6,201 at 10-100 RPKM, 988 at 100-1000 RPKM, and 228 exceed 1,000 RPKM (Fig. S5, Wang et al., 2019).

More than half of the genes in the 1-1000 RPKM regime harbour at least one CTCF site (Fig. S5A). However, intra-genic CTCF site orientation is not specific since the 4 orientation classes (Fig. 2C) are similarly represented (Fig. S5B). Stratification of CTCF sites into 4 quartiles (see Methods) shows that slightly more 1^st^ than 2^nd^, 3^rd^ or 4^th^ quartile CTCF sites reside inside transcription units (Fig. S5C). The highest proportion of genes overlapping TAD boundaries concerns the 6,201 genes expressed between 10 and 100 RPKM, including many boundaries observed in seven primary blood cell types (Fig. 3B, Fig. S5D).

The prevalence of CTCF sites inside transcribed genes and the prevalence of genes that overlap a conserved TAD boundary indicates that elongating RNA polymerase II is not inhibited by occupied CTCF sites, even those making-up TAD boundaries. We speculate that, beyond helping to compact the genome, the intragenic CTCF sites may also facilitate mechanistic aspects of gene expression control such as bringing together gene promoters and enhancers (eg; Fig. S1D), or marking alternative exon usage (Ruiz-Velasco et al., 2017). Thus, ‘intragenic’ CTCF sites are likely to contribute to the formation of CTCF-determined chromosome topology together with ‘intergenic’ CTCF sites.

## Discussion

In the budding yeast lower eukaryote model, inter-nucleosome-crosslinking studies revealed the existence of 3,061 boundaries that flank one to five genes and are spaced by an average of 4 ± 2 kb of DNA. These boundaries delimit many instances of small ‘loops’ connecting nucleosomes within ‘chromosomally interacting domains’. Interestingly, these yeast boundaries are enriched in strong promoters, such as those coding for ribosomal proteins (Hsieh et al., 2015).

Also in budding yeast, Cohesin is known to accumulate at intergenic regions located between convergent genes, indicating that transcribing RNA polymerase II enzymes can ‘push’ topologically trapped Cohesin complexes up to their polyadenylation sites (Glynn et al., 2004; Lengronne et al., 2004). Yeast Cohesin has been visualised at the single molecule level in vitro; DNA translocases can move Cohesin rings along DNA and a single nucleosome acts as a permeable barrier for Cohesin diffusion (Stigler et al., 2016).

In human cells, Cohesin complexes only accumulate at RNA polymerase II poly-adenylation sites when CTCF is absent (Busslinger et al., 2017). Otherwise, in the presence of CTCF, Cohesin rings embrace two convergent CTCF sites in more than 50% of the cases (Rao et al., 2014). Thus, Cohesin ring localisation can be a by-product of transcription, but CTCF-mediated signals appear to dominate the chromosomal loop landscape through a hypothetical Cohesin-based loop extrusions system that does not require transcribing RNA polymerase II enzymes. We investigated this theoretical framework in great detail at the *FKBP5* locus by strategically deploying 15 one-to-all 4C viewpoints and concluded that, also at this locus, convergent CTCF sites do tend to form loop anchors.

Two types of 4C viewpoints emerged from the analysis of long genes; Type I highlight the interior of convergent CTCF loops and Type II highlight loop edges. The juxtaposition of two loop edges revealed by 4C appears to be a boundary. Notably, boundaries harboured one or more divergent CTCF pairs.

To elucidate the relationship between CTCF sites, TAD boundaries and TAD centres, we introduced a triplet CTCF site classification and we combined this with a gradient of boundary identity based on blood cell TAD boundary conservation. Every human CTCF site was classified as a function of its own orientation and that of its two neighbouring CTCF sites’ orientations. Same (>>>, <<<) and convergent + divergent (<><, ><>) CTCF triplets do not change the polarity of the CTCF system, in that the CTCF site’s left and right neighbours have the same orientation. In the collection of 60,079 human CTCF sites we used here (Rao et al., 2014), there are 16,017 same CTCF sites and 14,369 convergent + divergent CTCF sites, indicating an evolutionary gain of same triplets in our genome. By contrast, there are as many convergent (>><, ><<) as divergent (<>>, <<>) triplet sites on any one piece of DNA, because convergent and divergent sites must operate in alternating fashion, respectively to change the orientation of the CTCF system from right to left-pointing and from left to right-pointing. More precisely, there are 15,305 convergent and 15,304 divergent sites in the collection of 60,079 human CTCF sites, when taking the 46 telomeres and 23 centromeres into account. It is probably this mathematical property of obligate alternation of convergent and divergent triplet, or tetra-plet (>><<, <<>>), CTCF sites that underlies the logic of chromosome loop network organisation (Fig. 6C).

At the hand of the triplet CTCF site classification we demonstrate that TAD boundaries are specifically enriched only in divergent CTCF triplets, as had been proposed for the Six homeobox gene cluster (Gomez-Marin et al., 2015). These TAD boundaries are defined as sites of abrupt direction change in the polarity of DNA-DNA interactions whereby the DI switches from a negative (towards the left) sign to a positive (towards the right) sign. Therefore, somewhere before the next boundary, the DI sign must switch from positive to negative. The (+) to (-) DI sign inversion loci delimit TAD halves. We show here that (+) to (-) DI index sign changes occur at loci that are depleted of same, divergent and convergent + divergent, but not of convergent CTCF sites.

Generally, a TAD can therefore be represented as <>(>…>)(<…<)<> with the boundaries harbouring divergent CTCF sites and the interior of the TAD harbouring a variable number of convergent sites. In support of this, we found that arrays of four or more CTCF sites display average directionality index sign biases that are consistent with this model. However, many left halves of TADs do harbour left-pointing arrays of same CTCF sites and vice versa, many right halves harbour right-pointing CTCF arrays. We speculate that this reflects evolutionary pressure to preserve locus-specific topologies that assist long-distance gene expression control via distal enhancers, as well as the evolutionary history of chromosomes during which inversions, deletions, duplications and translocations invariably took place.

We explored the relation between the CTCF topology of chromosomes and gene expression control at the hand of long-range epigenomic signalling by GR in monocytes and macrophages. Long-distance *cis*-activation of H3K27ac accumulation instigated by DNA-bound GR very rarely crosses a macrophage TAD boundary (D’Ippolito et al., 2018; Wang et al., 2019). Here we show that epigenomic *cis*-activation by GR is largely restricted to blood cell TAD halves. At *FKBP5*, the primary CTCF loop brings together the GR responsive enhancer in intron 5 and the upstream enhancer which are 118 kb apart, simply because they respectively flank the convergent CTCF20(>) and CTCF27/28(<<) sites that form the primary CTCF loop of the *FKPB5* TAD (Fig. 1, S1D). Such regulatory consequences are likely to be locus-specific and idiosyncratic (Ruiz-Velasco et al., 2017). Indeed, it is fascinating that 5,886 gene promoters are equipped with a functional CTCF site, such as CTCF11 at *PPARD* and CTCF16 at *RPL10A*. These may play key roles by insulating the DNA segments and coordinating aspects of gene expression control in the context of locally hard-wired chromatin loop networks.

Since we studied the TAD boundaries of seven primary white blood cell types, the question arose as to whether cell-type-specific boundaries exist. The answer is probably yes, since there are small subsets of boundaries that were only detected in two or three cell types, but this appears to be an exception rather than a rule. Similarly, Snyder and co-workers characterized about 16,000 chromatin loops in THP-1 cells, of which only 217 changed significantly upon induction of the monocyte to macrophage differentiation program by phorbol esters (Phanstiel et al., 2017). In line with the cell type-specific TAD hypothesis, ES-cells, have been reported to harbour ES cell-specific loops (Dixon et al., 2015).

To gauge the extent of cell type specificity, it may be fruitful to recoup CTCF motif CpG dinucleotide DNA methylation dynamics with boundary function, since cytosine methylation is known to lead to differential CTCF binding at epigenetically imprinted loci (Bell and Felsenfeld, 2000; Hark et al., 2000; Hashimoto et al., 2017; Parelho et al., 2008; Sun et al., 2013; Teif et al., 2012; Wendt et al., 2008; Wiehle et al., 2019).

The present conciliation of CTCF site orientation and TAD structure has deep implications for the further discovery, study and engineering of TADs and their boundaries. We expect that the mechanistic rules we report here will also be useful to conduct TAD and TAD boundary machine learning that should eventually yield a comprehensive atlas of human chromosome TADs, sub-TADs and boundaries.

## Acknowledgments

We wish to thank the members of the Molecular Biology department of the RIMLS and of the GeCo research group members at the Department of Electronics, Information and Bioengineering of Politecnico di Milano for their help in the course of this research. LN and SC are supported by Advanced ERC Grant 693174 “data-driven Genomic Computing” (GeCo).

## Author contributions

Conceptualisation: CL, LN, SC. 4Cseq analysis: FM, CW, LG, RO, PH. Bioinformatic analysis: LN. Paper writing: LN and CL.

## STAR METHODS

### Key resources table

| Reagent or Resource | Source | Identifier |
| --- | --- | --- |
| Ficoll | GE Healthcare | GE17-1440-02 |
| Percoll | Sigma-Aldrich | P1644-1L |
| DpnII | NEB | R0543M |
| NIaIII | NEB | R0125L |
| Bfal | NEB | R0568L |
| T4 ligase | NEB | M0202M |
| T4 ligase buffer | NEB | B0202S |
| Expand long template PCR system | Roche | 11681842001 |
| RPMI 1640 Medium, GlutaMAX | Thermo Fisher | 61870036 |
| L-glutamine | Thermo Fisher | 25030081 |
| Sodium Pyruvate | Thermo Fisher | 11360070 |
| dNTP set 100 mM | Thermo Fisher | 10297-018 |
| Sodium Acetate (3M) | Thermo Fisher | AM9740 |
| Glycogen | Thermo Fisher | 10814-010 |
| **Critical Commercial Assays** |  |  |
| CD3 MicroBeads, human | Miltenyi Biotec | 130-050-101 |
| CD19 MicroBeads, human | Miltenyi Biotec | 130-050-301 |
| CD56 MicroBeads, human | Miltenyi Biotec | 130-050-401 |
| KAPA library preparation kit | Kapa Biosystems | KK8400 |
| **Deposited Data** |  |  |
| GM12878 CTCF binding sites with orientation | (Rao et al., 2014) | Obtained from authors |
| GM12878 HiCCUPS long-range loops | (Rao et al., 2014) | ftp://ftp.ncbi.nlm.nih.gov/geo/series/GSE63nnn/GSE63525/suppl/GSE63525_GM12878_primary%2Breplic |
|  |  | ate_HiCCUPS_looplist.txt.gz |
| CTCF Chip-seq experiments from ENCODE | (Encode Consortium, 2012) | http://genome.ucsc.edu/cgi-bin/hgFileUi?db=hg19&g=wgEncodeOpenChromChip |
| TAD definitions from blood cell types | (Javierre et al., 2016) | https://osf.io/u8tzp |
| PC-HiC interactions | (Javierre et al., 2016) | https://osf.io/u8tzp |
| Genomic Regulatory Blocks | (Harmston et al., 2017) | https://static-content.springer.com/esm/art%3A10.1038%2Fs41467-017-00524-5/MediaObjects/41467_2017_524_MOESM2_ESM.txt |
| FastQ files of 4Cseq experiments | This paper | GEO accession GSE136034 |
| **Software and Algorithms** |  |  |
| PyGMQL | (Nanni et al., 2018) | https://github.com/DEIB-GECO/PyGMQL |
| HOMER | (Heinz et al., 2010) | http://homer.salk.edu/homer/ |
| Bedtools | (Quinlan, 2014) | https://github.com/arq5x/bedtools2 |
| PyBedtools | (Dale et al., 2011) | https://github.com/daler/pybedtools |
| CTCF clustering algorithm | This paper | https://github.com/lucananni93/hic_reconcile_ctcf/ |
| Boundary consensus algorithm | This paper | https://github.com/lucananni93/hic_reconcile_ctcf/ |
| CTCF looping simulation | This paper | https://github.com/lucananni93/hic_reconcile_ctcf/ |
| Hi-C normalization | (Imakaev et al., 2012) | https://github.com/mirnylab/cooler |
| DI index computation and boundary calling | (Abdennur and Mirny, 2019) | https://github.com/mirnylab/cooltools |
| Hi-C visualization | (Durand et al., 2016b) | https://www.aidenlab.org/juicebox/ |

### Ethical statement

Sanquin blood bank donors gave written informed consent for epigenetic research (NVT0068.01). All methods were carried out in accordance with, and using protocols approved by, the Sanquin Ethical Advisory Council, as previously described (Saeed et al., 2014; Wang et al., 2019).

### Cell culture

Peripheral blood mononuclear cells (PBMCs) were isolated from buffy coats using a Ficoll density gradient. Subsequently, monocytes were separated on a Percoll density gradient. Monocytes were purified via a negative selection kit using MACS CD3, CD19 and CD56 beads. Purified monocytes were cultured in RPMI1640 medium, supplemented with 10 μg/mL gentamycin, 10 mM L-glutamine and 10 mM pyruvate, 10% human serum and Penicillin/Streptomycin.

HeLa adherent (human cervical cancer cells) were cultured in Dulbecco’s Modified Eagle’s Medium (DMEM), supplemented with 10% Fetal Bovine Serum (FBS) and Penicillin/Streptomycin. THP-1 cells (human AML-derived monocytes) were grown in Roswell Park Memorial Institute (RPMI) 1640 medium supplemented with 10% FBS and Penicillin/Streptomycin. For glucocorticoid stimulation, cells were treated 100 nM Triamcinolone Acetonide (TA) or equal volume of DMSO for 4 hours.

### 4C Library Preparation

Circularised chromosome conformation capture (4C) assays were performed using a modified published approach (Kuznetsova et al., 2015; van de Werken et al., 2012; Wang et al., 2019). Ten million cells were crosslinked in 1% formaldehyde for 10 minutes and quenched with glycine. Cells were lysed in 30 ml lysis buffer (50 mM Tris pH 7.5, 150 mM NaCl, 0.5% NP-40, 1% Triton, 5 mM EDTA and 1X protease inhibitor cocktail (Roche)) at 4°C for 30 min. Isolated nuclei were digested using DpnII (NEB) followed by overnight ligation at 16°C using 50U T4 DNA ligase (NEB). The ligated library was de-crosslinked by incubating with proteinase K at 65°C and an RNase A treatment at 37°C was performed to remove RNA. Subsequently, second round of digestion using NlaIII or BfaI was performed. Digested DNA was ligated overnight and purified using a Qiagen Quick PCR purification kit. The 4C libraries were amplified with viewpoint-specific primers by inverse PCR. For each viewpoint, 8 PCR reaction products were pooled to enhance library complexity. The 4C PCR products were purified using Qiagen Quick PCR purification Kit.

### Next generation sequencing

4Cseq library preparation was performed using the KAPA hyperprep kit (KAPA Biosystems) according to the manufacturer’s protocol. Briefly, 10 ng DNA was incubated with end-repair and A-tailing buffer and incubated for 30 minutes at 20 °C followed by 30 minutes at 65 °C. Subsequently, samples were ligated at 20 °C for 15 minutes. Post-ligation clean-up was performed using Agencourt AMPure XP beads. 0.8X bead volume was added to the DNA and incubated for 10 minutes at room temperature, followed by two 80% freshly-made ethanol washes. DNA was eluted from beads and amplified using KAPA HiFi Hotstart ReadyMix and Primer Mix. DNA was purified using AMPure XP purification system with double size selection method to selected DNA fragment of 300 to 1000 bp. DNA library fragment size was checked using a BioAnalyzer. Sequencing was performed on the Illumina NextSeq 500 platform.

### CTCF binding sites dataset

For our analysis we used the CTCF binding sites from GM12878 (Rao et al., 2014) together with their motif orientation. We refer to a motif as ‘right’ (>) when it is present on the forward strand of the chromosome, while we call it ‘left (<) it is on the reverse strand.

### Assigning scores to CTCF binding sites

We downloaded the 33 ENCODE CTCF Narrow Peak tracks (Table S1) from the UCSC Browser^1^. For each CTCF binding site we then associate its enrichment signal for each of the Chip-seq tracks (using the *map* operation of PyGMQL (Masseroli et al., 2019). Before aggregating the 33 signal values for every CTCF binding site, we assessed the value distribution of every CTCF Chip-seq experiment and found heterogeneous distributions across cell lines, lineages and laboratories. To use comparable input values before computing the Chip-seq scores for the binding sites, we performed a quantile normalization across all the experiments. For each of the CTCF binding sites, we then combined the signal of the 33 ENCODE tracks by summing up their values; we call this Chip-seq score (*s*_*chipseq*_).

HOMER motif calling software was used to assess motif quality at each binding site; we call the motif matching result from the software motif score (*s*_*motif*_).

Since motif and Chip-seq scores have different distributions, the former having a normal-like distribution while the latter has a Poisson-like distribution, we normalize both values by taking their rank and multiply them to take into account both contributions. Therefore, the aggregated rank score of a CTCF binding site is

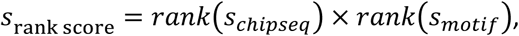

where `rank` returns a position in the ordered sequence of scores.

### CTCF spatial distribution analysis

We designed a clustering procedure of CTCF binding sites based on their distance. Given a clustering window *w*, each cluster is composed of a set of adjacent CTCF sites whose distance is less than or equal to *w* and they are not interrupted by a centromere. Then, for each cluster, we extract all mono-plets, di-plets, tri-plets and tetra-plets; the distance between each pair of CTCF sites composing one of these patterns will therefore be less than or equal to *w* (Fig. S2H). This clustering procedures identifies, at varying *w*, how orientation patterns change density along the genome. Notice that if a pattern is identified using a clustering window *w*, it will be identified also by all windows *w*^′^ ≥ *w*. When *w* → ∞ we end up with one cluster for each chromosome arm and with the complete set of *n-*plets.

We then perform, for each pattern *p* (composed by *n* binding sites) and each clustering window *w*, a statistical evaluation asking if *p* was over-represented or under-represented with respect to a uniform distribution of patterns with the same number of sites. This is done by performing a binomial test with null distribution 1/2^n^ (permutations of *n* symbols with 2 possible values), number of successes equal to the number of instances of *p* at window *w* and total number of trials equal to the total number of *n-*plets found at window *w*. We evaluated both the positive (over-representation) and negative (under-representation) one-sided tests for each *(p, w)* combination. In Fig. 2D and Fig. 2SA-F the p-values obtained by this procedure are displayed in red-scale for positive one-side test and blue-scale for negative one-side test. Formally, the value displayed in the figures is

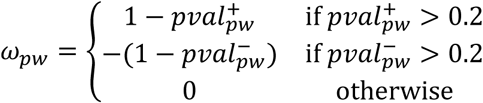

where 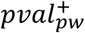 and 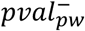 are respectively the right and left one-sided binomial test p-values for pattern *p* at clustering window *w*. In this way, we display both statistical tests using a divergent colour map with significant values at the extremes.

We performed the above procedure initially with the complete set of CTCF sites, and then by firstly selecting only CTCF sites with a *s*_rank score_ in ranges defined by the quartiles of rank score distribution, which are specified in Fig. S2B.

### Assigning a class to CTCF binding sites

We assigned each CTCF binding site to a triplet class depending on its two adjacent sites. There are therefore eight different possible patterns, which we then name following the classification of CTCF triplets shown in Fig. 2C. Notice that the classification concerns the central CTCF site of the triplet, and that this is independent of the distances between CTCF sites forming the triplet.

### Boundary consensus algorithm

We developed a novel algorithm to find the consensus region across several TAD boundary datasets. The algorithm takes as input a set of boundary datasets in the form of genomic regions and a *detection window* (*w*), which determines the distance up to which two *boundary positions* (the start or stop position of a TAD boundary) can be considered to belong to the same cluster. In our study, given the minimum Hi-C resolution adopted by (Javierre et al., 2016) of 25kb, we imposed *w=25kb*. The algorithm builds an adjacency matrix *A* where every row and column represent a boundary position across the whole set of experiments. A cell of *A* is equal to one if its two positions have a distance less or equal than *w*. This makes *A* a symmetric matrix. We then apply the Louvain modularity (Subelj and Bajec, 2011) community detection algorithm to *A* and find clusters of boundary positions. The *consensus region* of a cluster is the genomic region between the most upstream and most downstream boundary position. The centre of the computed boundary is halfway. The *boundary conservation score* is the number of cell types from which the region derives (Fig. 3A).

### GM12878 boundaries call

We downloaded the GM12878 ‘primary + replicate combined’ *.hic* file from GSE63525. We used the Cooler software (Abdennur and Mirny, 2019) for data processing and analysis. Coherently with (Javierre et al., 2016) we performed iterative correction of the contact matrix (Imakaev et al., 2012) and then calculated Directionality Index (Dixon et al., 2012) at a resolution of 25 kb. Boundaries were called at peaks of insulation score with the cooltools suite (`cooltools diamond-insulation` command).

### Intersecting GM12878 boundaries with consensus boundaries

Since the length of the consensus boundaries is variable and dependent on their conservation score, we removed length biases by taking the centre of each boundary 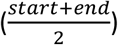 using BedTools (Quinlan, 2014) and extended it 25 kb to the left and to the right. We then counted how many boundaries intersect the consensus boundaries and stratified the result by conservation score (Fig. 3E).

### Relationship between PC-HiC interactions and consensus boundaries

We downloaded the PC-HiC interactions with CHICAGO score greater than 5 from (Javierre et al., 2016). Following the strategy of (Schoenfelder et al., 2015), we computed the percentage of promoter-genome interactions which span a boundary. We then computed the same metric using the boundary set shifted in steps of 10 kb to the left and to the right up to 400 kb (Fig. 3F). We stratified the resulting curves based on the different sets of boundaries built progressively by filtering on the conservation score. Therefore the *s* ≥ 1 set contains all the boundaries detected by the consensus algorithm, *s* ≥ 2 contains all boundaries seen in *at least* two cell datasets, up to *s* ≥ 7.

### Relationship between GRBs and consensus boundaries

We downloaded the dataset of GRBs from Harmston 2017 (Harmston et al., 2017). To determine how our consensus boundaries relate with these highly conserved regions, we ordered GRBs based on their length and aligned them on their centre. We then plotted the position of *s* ≥ 2 consensus boundaries in a 5 Mb region around them (Fig. 3G). The results shows enrichment of boundaries at the borders of GRBs and depletion inside, demonstrating high concordance between the datasets.

### Enrichment analysis of CTCF patterns at consensus boundaries

We counted how many boundaries harbour zero, one or more CTCF binding sites within less than 25 kb of their limits. The results were stratified by conservation score (Fig. 4A). We associated to each CTCF binding site a unique class depending on its two flanking CTCF sites using the same nomenclature introduced for 3-plets in Fig. 2C, where the pattern is for the middle site, as determined by the orientation of both its neighbours. We then took the centres of all consensus boundaries and looked at the region 250 kb upstream and downstream and computed the average number of CTCFs (Fig. 4B) and then for each orientation (Fig. 4C) and class (Fig. 4D) in windows of 5 kb.

### Enrichment analysis of CTCF patterns at TADs

We investigated the average enrichment of CTCF sites (Fig. 5A), their orientation (Fig. 5B) and their classes (Fig. 5C) on TADs by dividing each of them in 100 bins and counting how many binding sites fall inside each of them. Since TADs have variable size, the size of bins for each TAD is variable. We then aggregate all the bin values in a single 100-valued TAD vector representing the average enrichment. The x-axis in Fig. 5 can be seen as relative positions with zero representing the beginning and one the ending of TADs.

### Analysis of Directionality index and CTCF patterns in TADs

We took the DI index computed at 25 kb resolution on GM12878 and identified points of negative inversion (+) to (-) along the genome. While points of positive inversion (-) to (+) are an indication of sudden increase of insulation determining a boundary, the negative inversions are smoother and tend to localize towards the middle of TADs. We then looked at the region 250 kb upstream and downstream to each DI index negative inversion point and computed the average number of CTCFs (Fig. 5D) and then of each orientation (Fig. 5E) and class (Fig. 5F) in windows of 5 kb. We then extracted from the whole dataset of CTCF binding sites, all the sets of same-oriented motifs of every length. We looked at the average DI index in the regions covered by these sequences. We observed a strong predominance of positive DI values for Forward patterns and negative DI values for Reverse patterns, and this correspondence sharpens with the length of the pattern (Fig. 5G).

Each TAD can be seen as the composition of two regions; the first starts at the point of positive inversion and ends at the one of negative inversion (*positive DI*), the second starts at the point of negative inversion and ends at the one of positive inversion (*negative DI*). We took all positive and negative DI regions and reported the presence of forward and reverse CTCF binding sites ±1 *Mb* region around them. Finally, we ordered the regions by size (Fig. 5H).

### Glucocorticoid induction analysis in DI index delimited regions

We downloaded the dataset of robust glucocorticoid peaks and the epigenetic H3K27ac response (Wang et al., 2019). We then took the previously extracted positive DI and negative DI regions and selected only those bearing at least one GR peak inside, but whose four adjacent regions do not have any GR peak. We then counted the number of up-regulated and down-regulated acetylation peaks or genes in all the selected regions and their neighbours (Fig. S4).

### CTCF looping extrusion simulations

To simulate loop extrusion, we first assign to each CTCF binding site a *permeability* score, which is a number between zero and one describing the probability of the extrusion complex to ignore a correctly oriented CTCF site during the pulling phase.

We first designed a ‘full-stop’ model in which all the CTCF sites have a permeability score of zero, meaning that the extrusion complex will always stop pulling DNA when it encounters a correctly oriented CTCF motif. We then defined a set of models based respectively on ChIPseq, motif and rank score. For each CTCF site *c*, its permeability score in these models is

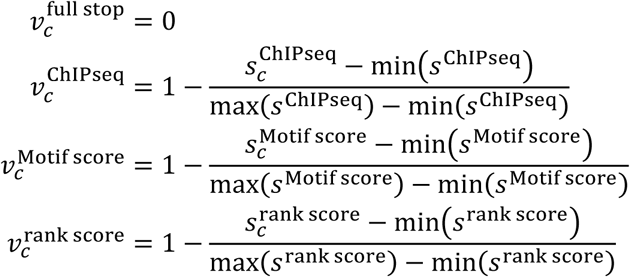

For each simulation epoch, we execute an extrusion event in which the hypothetical extrusion complex binds one time between two adjacent CTCF binding sites. Our simulations are therefore independent of CTCF density fluctuations along the genome. The hypothetical extrusion complex pulls DNA on its right and left at the same time and independently stops extrusion on each side when the following two conditions apply: firstly, the complex has reached a correctly oriented CTCF site and, secondly, a randomly generated number between 0 and 1 is lower than the permeability score of this CTCF site. A loop can be generated multiple times in the same epoch and across epochs. In this way we can model both the statistical variability of loop formation due to CTCF permeability and the geometrical constraints due to CTCF orientation spatial patterns. In our experiments we fixed the number of epochs to 100. To speed-up the execution we parallelized each simulation epoch.

### HiCCUPs comparison with loop simulations

We downloaded long-range loops extracted by the HiCCUPS algorithm from (Rao et al., 2014). We then computed the percentage of HiCCUPS loop (recall) that are recovered by the various models as a function of their occurrence in the simulations. Precision-recall curves of the indicated models were used to investigate the percentage of simulated loops that are present in the HiCCUPS collection as a function of recall.

---

**Fig. S1:**
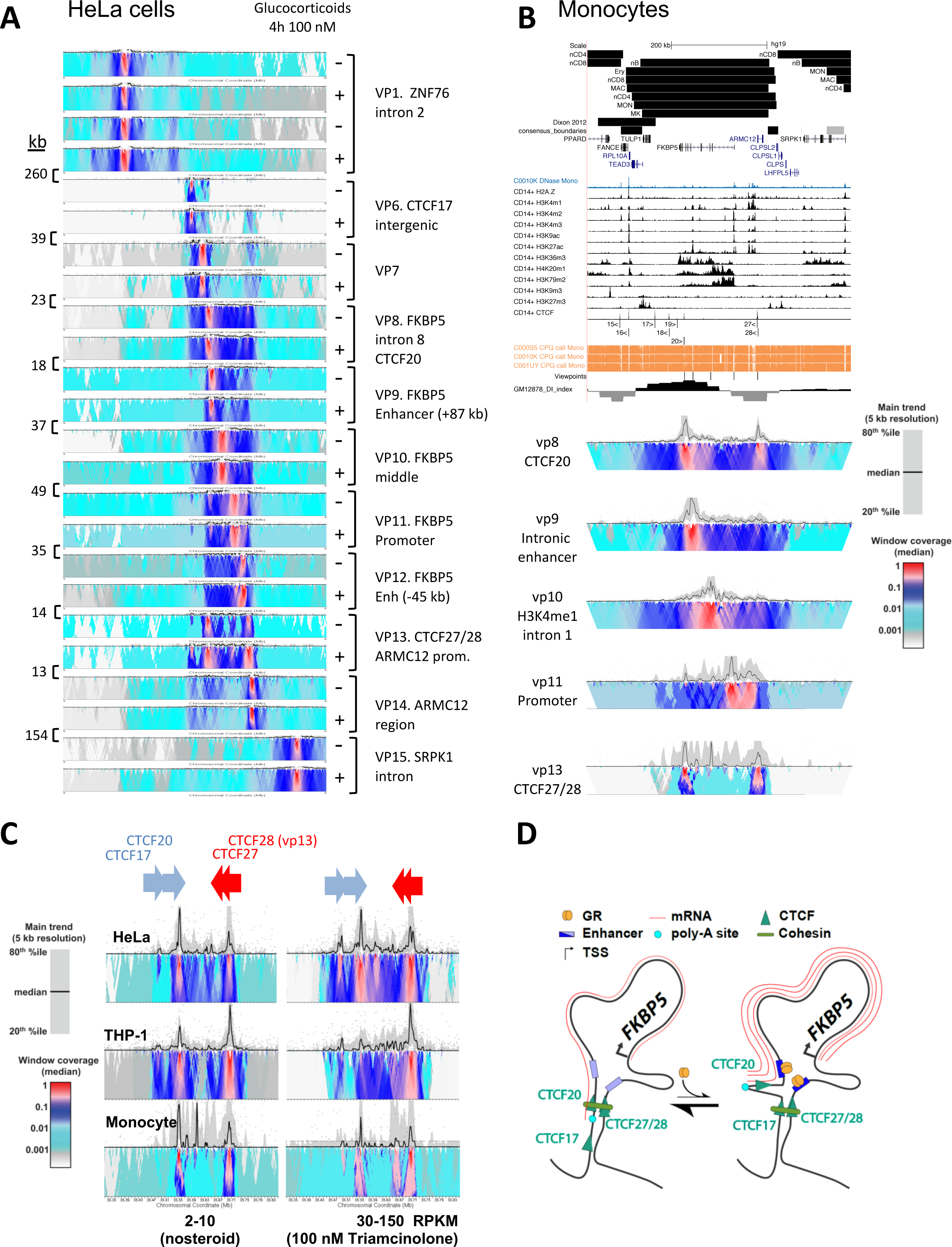
Transcription rate effects on *FKBP5* TAD structure. A Eleven viewpoints from HeLa cells treated with 100 nM triamcinolone acetonide (TA) or not are rendered by the 4C-seq pipeline (van de Werken et al., 2012). The distance between viewpoints is indicated (kb) and the viewpoints are named at the hand of close genomic feature of interest. B As for Figure 1, Javierre blood cell TADs are shown, above the Dixon 2012 (Dixon et al., 2012) and s1-s7 conserved consensus boundaries (present work), followed by Blueprint DNase accessibility in primary monocytes (Saeed et al., 2014), and 11 histone ChIPseq tracks as well as the CTCF ChIPseq track from ENCODE CD14+ cell UCSC genome browser tracks (Ram et al., 2011). Five viewpoints locations are shown above the Blueprint DNA methylation tracks for primary monocytes (bioRxiv, doi.org/10.1101/237784) above the GM12878 DI index. Below, viepwoints 8, 9, 10, 11 and 13 are shown for healthy blood donor primary human monocytes. C Viewpoint 13 was deployed in 3 human cell types treated or not for 4 hours with 100 nM triamcinolone acetonide. CTCF17, 20, 27 and 28 are indicated. D An ‘artist rendering’ of the 2D model of the *FKBP5* TAD architecture under low (left) and high (right) transcription regimes.

**Fig. S2:**
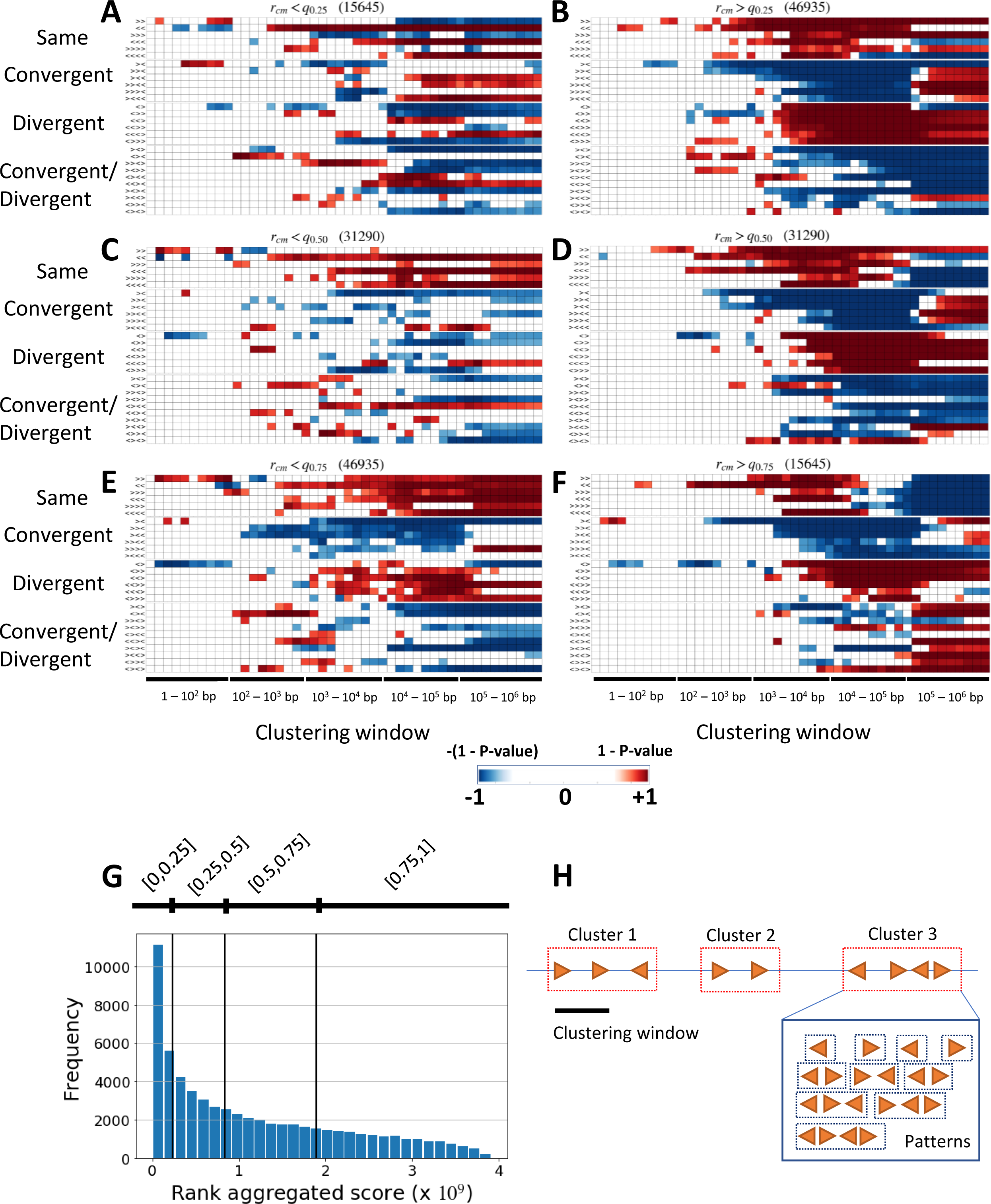
Stability of CTCF spatial patterns. A-F Statistical assessment of over- and under-represented CTCF patterns (see Fig.2d) by first removing CTCF binding sites in the low (a, c, e) and high (b, d, f) quartiles of the rank score distribution. On top of each heatmap, the quartile interval of CTCF sites kept during the analysis is reported together with their number. G Rank score distribution along the whole set of CTCF binding sites. Quartiles are highlighted by black vertical lines H Schematic representation of the clustering and pattern finding process.

**Fig. S3:**
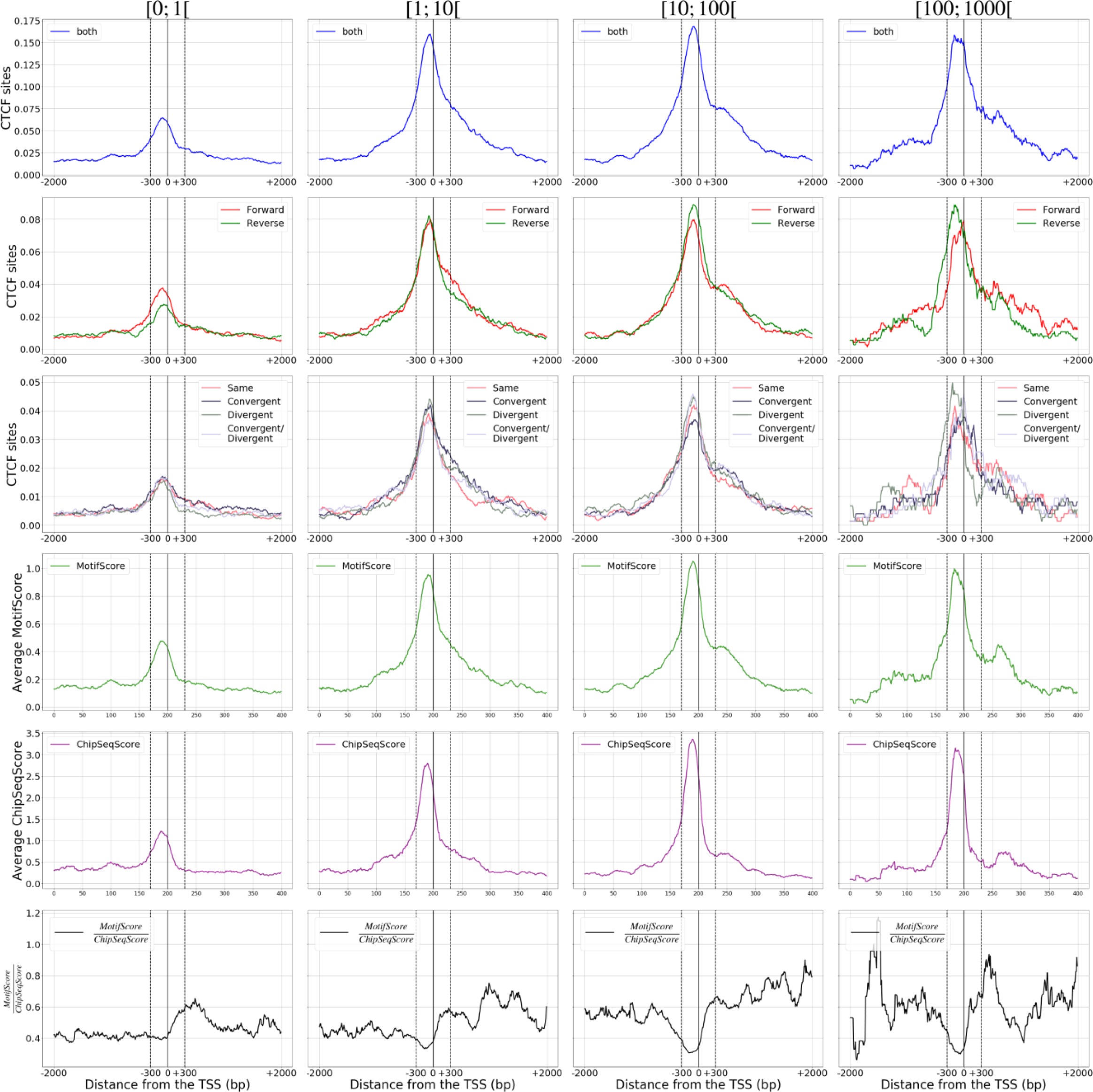
CTCF sites at human gene promoters. Promoters were stratified on their expression level in primary human monocytes (Wang et al., 2019). The number, direction, orientation class, motif scores, average ChIPseq scores and the ratio of Motif score to ChIPseq score are plotted.

**Fig. S4:**
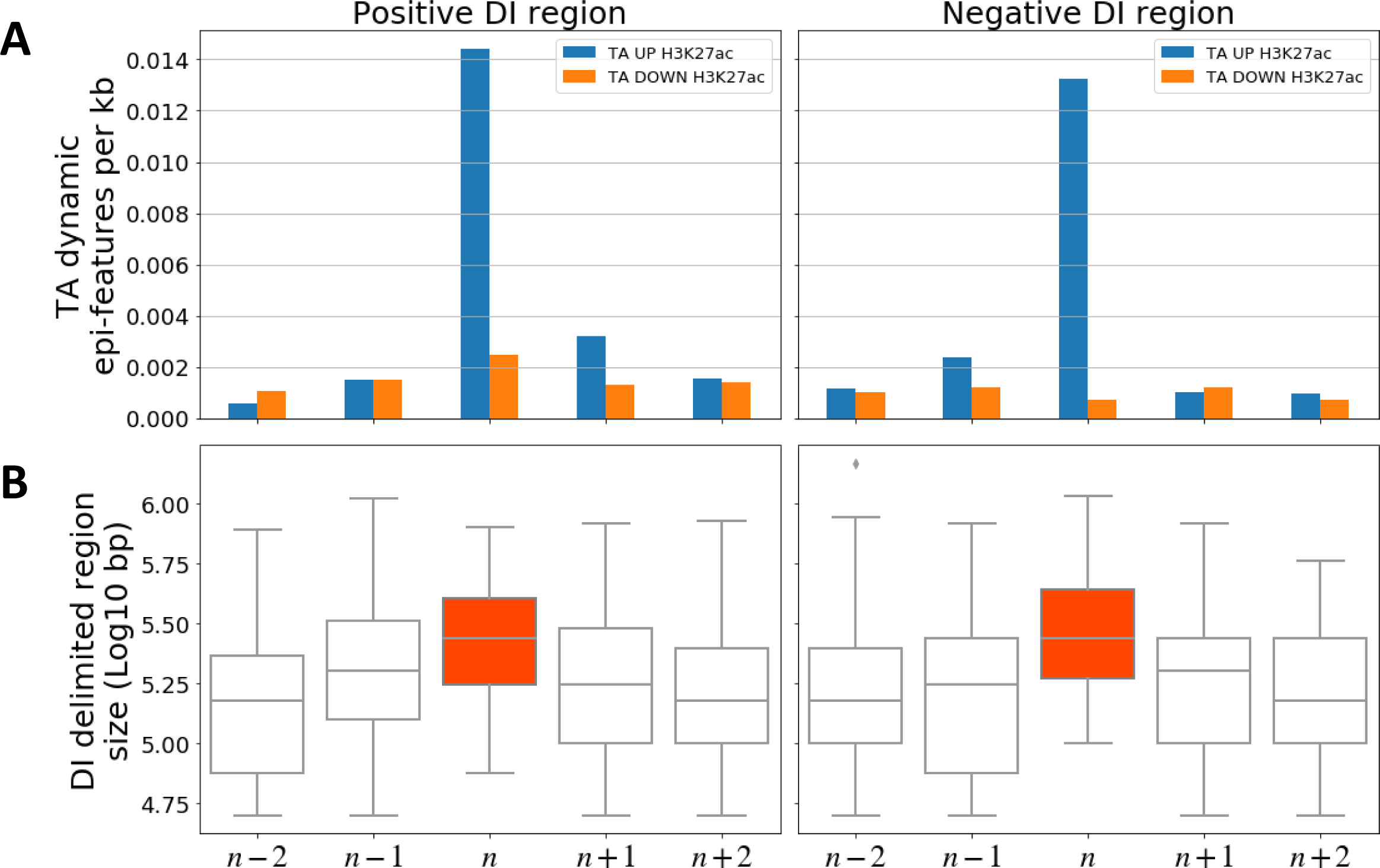
Glucocorticoid epigenomic signalling is conforms to half-TAD dimensions. A Epigenetic features (H3K27ac) induced (blue) and repressed (orange) by glucocorticoids in primary human monocytes projected on sets of five adjacent DI regions. The central region bears at least one GR ChIPseq site, the others do not. The results are stratified by DI index sign. B Size distributions of the five adjacent DI region sets.

**Fig. S5:**
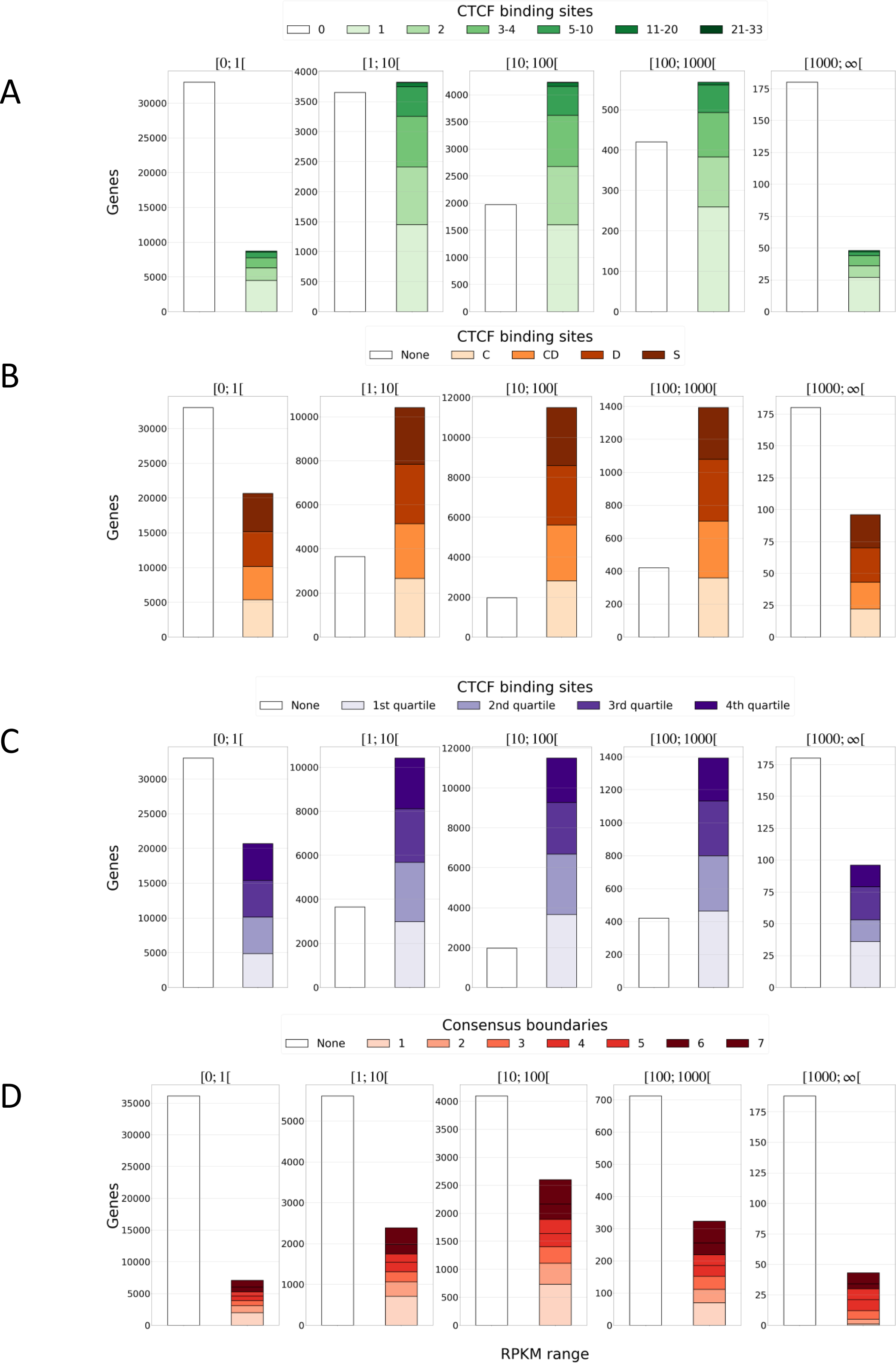
Intragenic TAD boundaries and CTCF site orientations. A Abundance of intragenic CTCF sites, ranging from none to 33, stratified as a function of gene expression level (Reads Per Kilobase per Million reads - RPKM) in healthy primary monocyte-derived macrophages (Wang et al., 2019). B Intragenic CTCF site orientation class (Fig. 2C) occurrence. The Y-axis thus represents the number of intragenic CTCF instances, rather than genes C Intragenic CTCF site strength, stratified as a function of their aggregate rank score (Fig. S2G), projected as in panel (B). D Number and conservation score of TAD boundaries that overlap genes (Fig. 3B), projected as in panel (A).

**Tab. S1:**
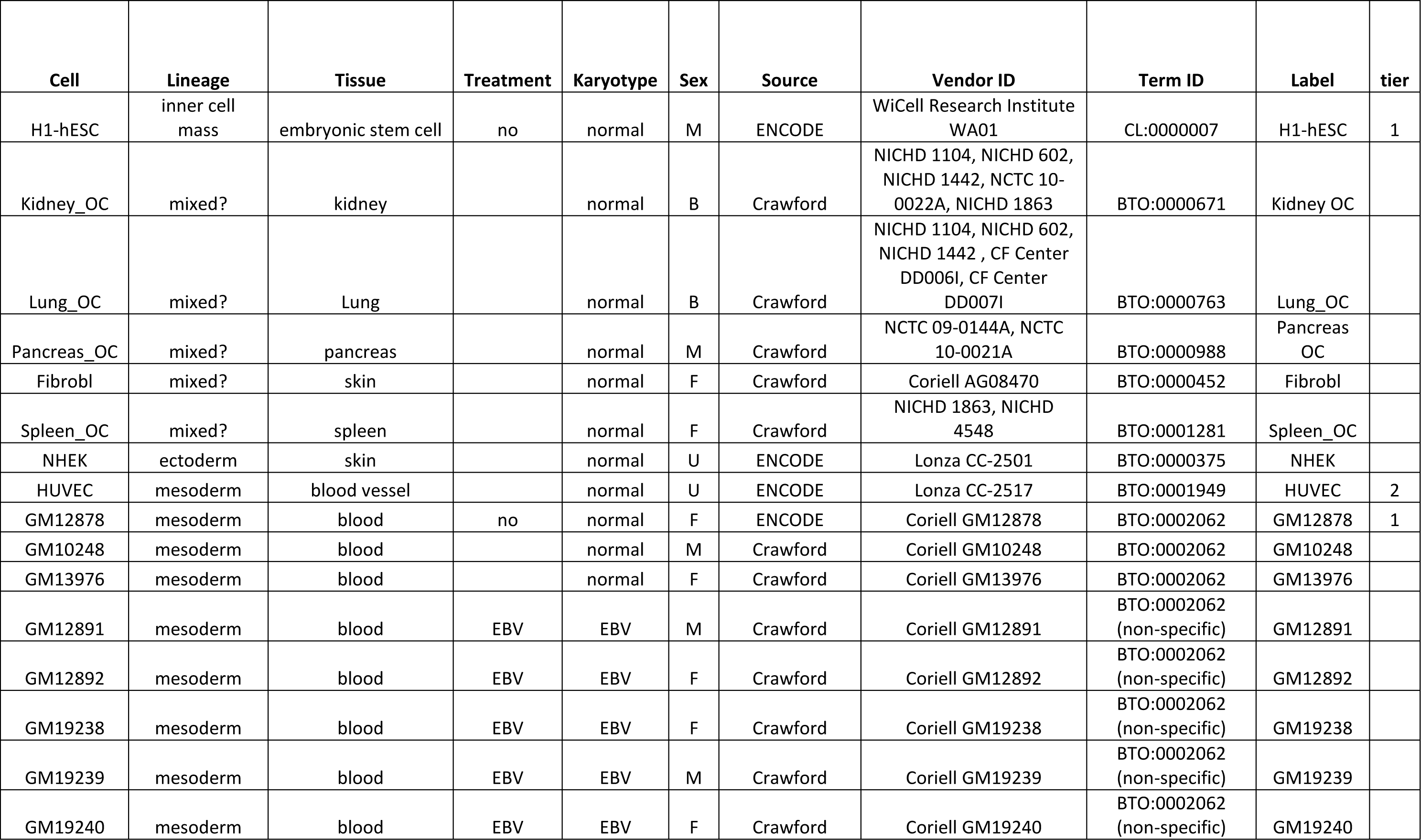
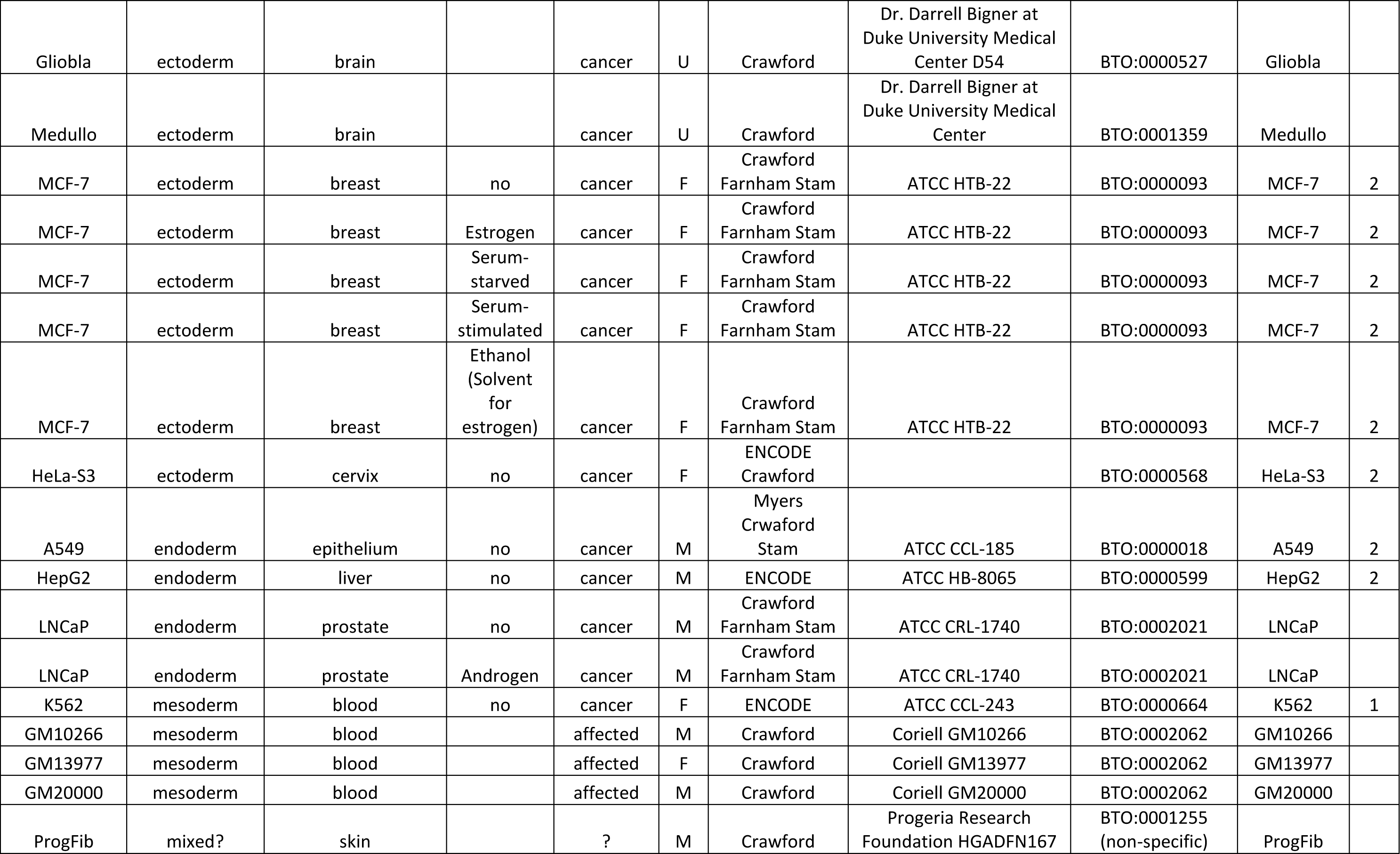
Metadata of 33 ENCODE samples of CTCF ChIPseq downloaded for this study

| Cell | Lineage | Tissue | Treatment | Karyotype | Sex | Source | Vendor ID | Term ID | Label | tier |
| --- | --- | --- | --- | --- | --- | --- | --- | --- | --- | --- |
| H1-hESC | inner cell mass | embryonic stem cell | no | normal | M | ENCODE | WiCell Research Institute<br>WA01 | CL:0000007 | H1-hESC | 1 |
| Kidney_OC | mixed? | kidney |  | normal | B | Crawford | NICHD 1104, NICHD 602,<br>NICHD 1442, NCTC 10-0022A, NICHD 1863 | BTO:0000671 | Kidney OC |  |
| Lung_OC | mixed? | Lung |  | normal | B | Crawford | NICHD 1104, NICHD 602,<br>NICHD 1442 , CF Center<br>DD006I, CF Center<br>DD007I | BTO:0000763 | Lung_OC |  |
| Pancreas_OC | mixed? | pancreas |  | normal | M | Crawford | NCTC 09-0144A, NCTC<br>10-0021A | BTO:0000988 | Pancreas<br>OC |  |
| Fibrobl | mixed? | skin |  | normal | F | Crawford | Coriell AG08470 | BTO:0000452 | Fibrobl |  |
| Spleen_OC | mixed? | spleen |  | normal | F | Crawford | NICHD 1863, NICHD<br>4548 | BTO:0001281 | Spleen_OC |  |
| NHEK | ectoderm | skin |  | normal | U | ENCODE | Lonza CC-2501 | BTO:0000375 | NHEK |  |
| HUVEC | mesoderm | blood vessel |  | normal | U | ENCODE | Lonza CC-2517 | BTO:0001949 | HUVEC | 2 |
| GM12878 | mesoderm | blood | no | normal | F | ENCODE | Coriell GM12878 | BTO:0002062 | GM12878 | 1 |
| GM10248 | mesoderm | blood |  | normal | M | Crawford | Coriell GM10248 | BTO:0002062 | GM10248 |  |
| GM13976 | mesoderm | blood |  | normal | F | Crawford | Coriell GM13976 | BTO:0002062 | GM13976 |  |
| GM12891 | mesoderm | blood | EBV | EBV | M | Crawford | Coriell GM12891 | BTO:0002062<br>(non-specific) | GM12891 |  |
| GM12892 | mesoderm | blood | EBV | EBV | F | Crawford | Coriell GM12892 | BTO:0002062<br>(non-specific) | GM12892 |  |
| GM19238 | mesoderm | blood | EBV | EBV | F | Crawford | Coriell GM19238 | BTO:0002062<br>(non-specific) | GM19238 |  |
| GM19239 | mesoderm | blood | EBV | EBV | M | Crawford | Coriell GM19239 | BTO:0002062<br>(non-specific) | GM19239 |  |
| GM19240 | mesoderm | blood | EBV | EBV | F | Crawford | Coriell GM19240 | BTO:0002062<br>(non-specific) | GM19240 |  |
| Gliobla | ectoderm | brain |  | cancer | U | Crawford | Dr. Darrell Bigner at<br>Duke University Medical<br>Center D54 | BTO:0000527 | Gliobla |  |
| Medullo | ectoderm | brain |  | cancer | U | Crawford | Dr. Darrell Bigner at<br>Duke University Medical<br>Center | BTO:0001359 | Medullo |  |
| MCF-7 | ectoderm | breast | no | cancer | F | Crawford<br>Farnham Stam | ATCC HTB-22 | BTO:0000093 | MCF-7 | 2 |
| MCF-7 | ectoderm | breast | Estrogen | cancer | F | Crawford<br>Farnham Stam | ATCC HTB-22 | BTO:0000093 | MCF-7 | 2 |
| MCF-7 | ectoderm | breast | Serum-<br>starved | cancer | F | Crawford<br>Farnham Stam | ATCC HTB-22 | BTO:0000093 | MCF-7 | 2 |
| MCF-7 | ectoderm | breast | Serum-<br>stimulated | cancer | F | Crawford<br>Farnham Stam | ATCC HTB-22 | BTO:0000093 | MCF-7 | 2 |
| MCF-7 | ectoderm | breast | Ethanol<br>(Solvent<br>for<br>estrogen) | cancer | F | Crawford<br>Farnham Stam | ATCC HTB-22 | BTO:0000093 | MCF-7 | 2 |
| HeLa-S3 | ectoderm | cervix | no | cancer | F | ENCODE<br>Crawford |  | BTO:0000568 | HeLa-S3 | 2 |
| A549 | endoderm | epithelium | no | cancer | M | Myers<br>Crawford<br>Stam | ATCC CCL-185 | BTO:0000018 | A549 | 2 |
| HepG2 | endoderm | liver | no | cancer | M | ENCODE | ATCC HB-8065 | BTO:0000599 | HepG2 | 2 |
| LNCaP | endoderm | prostate | no | cancer | M | Crawford<br>Farnham Stam | ATCC CRL-1740 | BTO:0002021 | LNCaP |  |
| LNCaP | endoderm | prostate | Androgen | cancer | M | Crawford<br>Farnham Stam | ATCC CRL-1740 | BTO:0002021 | LNCaP |  |
| K562 | mesoderm | blood | no | cancer | F | ENCODE | ATCC CCL-243 | BTO:0000664 | K562 | 1 |
| GM10266 | mesoderm | blood |  | affected | M | Crawford | Coriell GM10266 | BTO:0002062 | GM10266 |  |
| GM13977 | mesoderm | blood |  | affected | F | Crawford | Coriell GM13977 | BTO:0002062 | GM13977 |  |
| GM20000 | mesoderm | blood |  | affected | M | Crawford | Coriell GM20000 | BTO:0002062 | GM20000 |  |
| ProgFib | mixed? | skin |  | ? | M | Crawford | Progeria Research<br>Foundation HGADFN167 | BTO:0001255<br>(non-specific) | ProgFib |  |

## Footnotes

1 33 Encode CTCF tracks

